# CANVAS causing AAGGG repeat expansions cause tissue-specific reduction in RFC1 expression and increase sensitivity to DNA damage

**DOI:** 10.1101/2025.11.18.688292

**Authors:** Riccardo Curro, Natalia Dominik, Stefano Facchini, Ricardo Parolin Schnekenberg, Cecilia Perini, Riccardo Ronco, Bianca Rugginini, Arianna Ghia, Silvia Bione, Nadia Tagliaferri, Glenda P Grupelli, Simon A Lowe, Amy R Hicks, Elisa Vegezzi, Roberto Simone, Alessandro Bertini, Elena Abati, Roser Velasco, Maria Sereno, Gerardo Gutiérrez-Gutiérrez, Simone Thomas, Paola Alberti, Vikram Khurana, Johannes Attems, Claire Troakes, Emil K Gustavsson, Gabriele Lignani, Yichen Qiu, James N Sleigh, Arianna Tucci, Pietro Fratta, Adrian Isaacs, Yau M Lim, Zane Jaunmuktane, Sebastian Brandner, David L Bennett, Serge Przedborski, Puneet Opal, Ahmet Hoke, Sheng-Han Kuo, Mary M Reilly, Henry Houlden, Mina Ryten, Guido Cavaletti, Andreas A Argyriou, Jordi Bruna, Chiara Briani, Emmanuele Crespan, James EC Jepson, Andrea Cortese

## Abstract

Biallelic AAGGG expansions in *Replication Factor Complex Subunit 1* (*RFC1)* are associated with cerebellar ataxia, neuropathy, vestibular areflexia syndrome (CANVAS) and are increasingly recognised as a common cause of adult-onset ataxia and sensory neuropathy. However, the disease-causing mechanisms remain unclear. Here we leveraged *in vitro* assays, post-mortem brain tissue, patient-derived cell lines and a neuronal *Drosophila* model to demonstrate that AAGGG expansions are associated with tissue-specific reductions in the expression of *RFC1* transcript, along with impaired RFC1 function and increased sensitivity to DNA damage from platinum-based drugs. CRISPR/Cas9 excision of the AAGGG repeat and flanking AluSx3 element normalized *RFC1* expression in iPSC-derived neurons and rescued the DNA damage response, providing a framework for future therapeutic strategies. We also show that these biological findings are clinically relevant in heterozygous AAGGG expansion carriers, who display an increased risk and severity of neuropathy with platinum-based chemotherapy.

## INTRODUCTION

Biallelic AAGGG expansions (CCCTT in the coding strand) in the second intron of Replication Factor Complex Subunit 1 (RFC1) have been identified as the cause of cerebellar ataxia, neuropathy, vestibular areflexia syndrome (CANVAS) (Cortese et al. 2019; Rafehi et al. 2019), and are now recognised to be a common cause of late-onset ataxia (Traschütz et al. 2021; Vegezzi et al. 2024) and sensory neuropathy (Currò et al. 2021; Tagliapietra et al. 2021) in the ageing population.

To date, the mechanism by which AAGGG repeat expansions in *RFC1* cause neuronal dysfunction remains unclear. Initial studies did not show reduced *RFC1* expression or changes in *RFC1* mRNA processing in replicating cell lines and in post-mortem brain tissue from a CANVAS patient (Cortese et al. 2019; Gisatulin et al. 2020b). These findings were puzzling considering the recessive mode of inheritance of this condition.

More recent studies have suggested AAGGG repeat-derived RNA foci (Wada et al. 2024) and peptides (Maltby et al. 2024) as possible contributors to the disease pathogenesis. Indeed, patient-derived cortical motor neurons were shown to exhibit defects in neuronal development, reduced expression of synaptic genes, and diminished synaptic connectivity, which were partly rescued by CRISPR deletion of a single AAGGG expanded repeat, but not by *RFC1* supplementation (Maltby et al. 2024), implying that RFC1-independent mechanisms may drive CANVAS pathogenesis.

Yet, the identification of several CANVAS patients carrying nonsense variants in *RFC1* in *trans* with an expanded allele provide strong genetic evidence in support of a loss or reduced function model of the disease (Benkirane et al. 2022; Ronco et al. 2023). Similarly, point mutations *in trans* have been observed in Friedreich’s ataxia (FRDA), another intronic repeat expansion disorder, in which the loss of function of the encoded protein has been clearly demonstrated (Campuzano et al. 1996; Cossée et al. 1999; Bidichandani et al. 1998).

Moreover, clinical onset and severity of CANVAS appear to be more strongly dependent on the size of the smaller of the two expanded alleles, which may suggest an effect of the expanded repeat on residual gene expression and function (Currò et al. 2023), as also shown in Friedreich’s ataxia (Saccà et al. 2011; Plasterer et al. 2013).

RFC1 is the largest of the five subunits of the Replication Factor Complex (RFC), which is responsible for loading Proliferating Cell Nuclear Antigen (PCNA) onto the double-helix molecule during DNA replication and repair (Indiani and O’Donnell 2006; Mossi et al. 1997; Allen et al. 1998). Cells that are primarily affected in CANVAS, including cerebellar and sensory neurons, appear to be particularly sensitive to DNA damage, as showed by their primary involvement in other diseases linked to mutations in genes involved in DNA repair, including *ATM* causing ataxia-telangiectasia, *APTX* and *SETX* causing ataxia-oculomotor apraxia, *TDP1* causing spinocerebellar ataxia with axonal neuropathy-1, and several genes associated with xeroderma pigmentosum (reviewed in Kulkarni and Wilson 2008).

Prompted by these observations, we used cellular and animal models, post-mortem brain tissue, and clinical/genetic data from CANVAS patients, to study the effect of biallelic AAGGG expansions on neuronal *RFC1* expression and function. Additionally, given the high frequency of *RFC1* expansion carriers (8% of the European population) (Ibañez et al. 2024), we gathered evidence supporting a role for heterozygous AAGGG expansions as modifiers of the prevalence and severity of chemotherapy-induced peripheral neuropathy.

## RESULTS

### Pathogenic RFC1 repeats form stable parallel G-quadruplex structures

To investigate how *RFC1* expansion may affect cellular functions and contribute to neurological disease, we first analysed the repeat sequence for distinctive structural features. The DNA repeat sequence found in the template strand of *RFC1* contains evenly spaced runs of guanines and is predicted *in silico* to form G-quadruplexes (G4) (Dominik et al. 2023), which are secondary structures known to modulate transcription and stall the progression of DNA and RNA polymerase (Varshney et al. 2020).

To validate these *in silico* predictions, we compared 5’-(AAAAG)_8_-3’ (non-pathogenic) and 5’-(AAGGG)_8_-3’ (pathogenic) DNA sequences by circular dichroism spectroscopy (CD) in the presence KCl, NaCl, or LiCl –which promote (KCl) or impair (NaCl, LiCl) G4 stability (Kharel et al. 2020).

Consistent with(Varshney et al. 2020) previous studies (Abdi et al. 2023; Wang et al. 2024; Kudo et al. 2024; Hisey et al. 2024), the pathogenic 5’-(AAGGG) -3’ sequence preferentially forms a stable parallel G4 under physiologically relevant potassium conditions, while the non-pathogenic 5’-(AAAAG) -3’ sequence did not form G4 (**Fig. 1A and Supplementary Fig. 1A-B**). Principal component analysis (PCA) of CD spectra further highlighted this distinction: pathogenic AAGGG repeats clustered with G4 references (parallel in K , hybrid in Na and Li ), whereas AAAAG repeats are clearly separated from the G4 structural clusters, indicating that they do not adopt a canonical G4 structure (**Supplementary Fig. 1C**).

**Figure 1.**
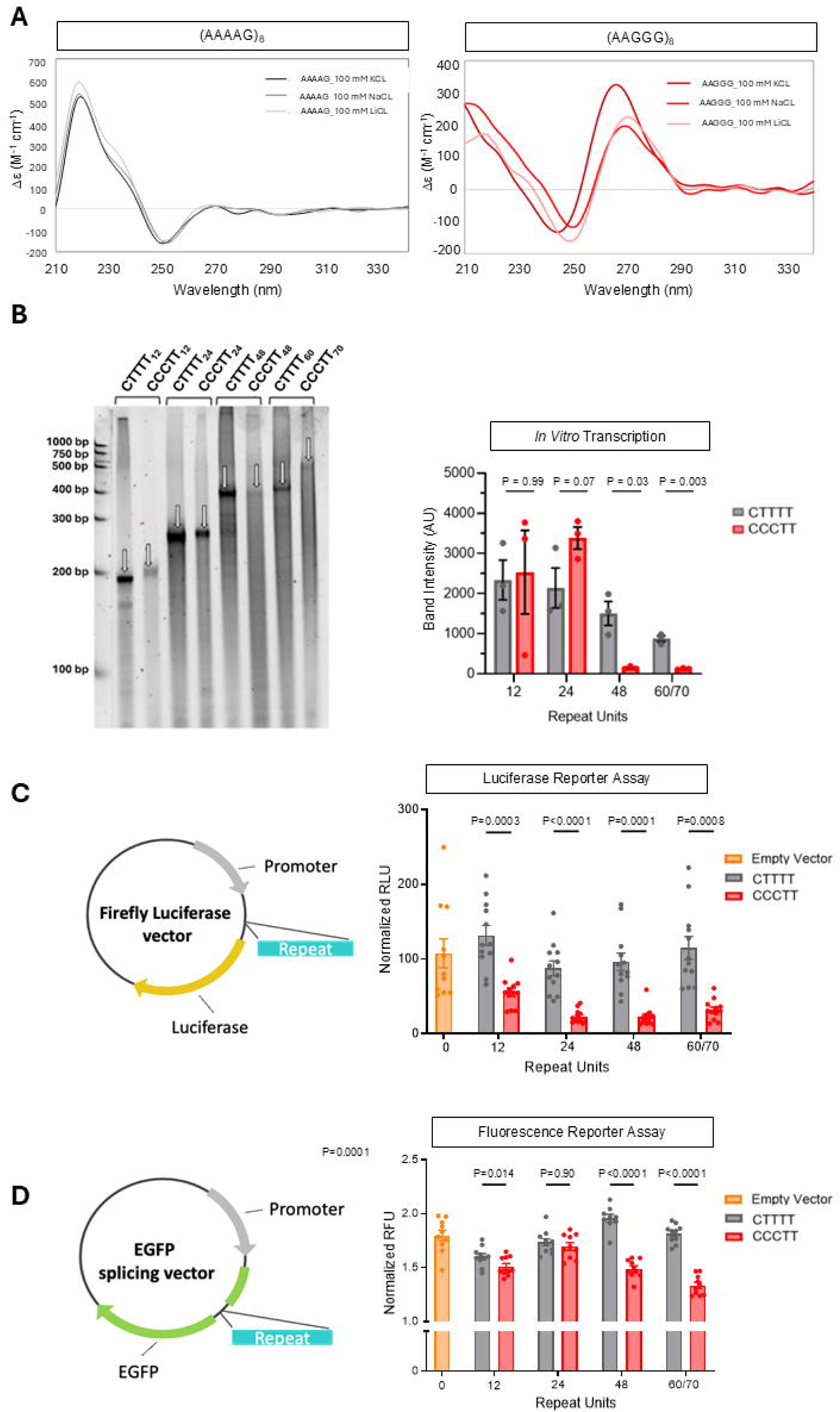
RFC1 repeat expansions form G quadruplexes and impair transcription and gene expression *in vitro models*. **A.** Circular dichroism spectra of 1 µM (A) 5’-(AAAAG)8-3’ and (B) 5’-(AAGGG)8-3’ measured in 10 mM lithium cacodylate buffer at pH 7.2, with 100 mM of KCl (black lines), NaCl (grey lines), or LiCl (light grey lines), maintained at 20°C. In 100 mM KCl 5’-(AAGGG) -3’ CD spectrum displayed a positive peak at 264 nm, a negative peak near 240 nm, and an additional positive peak around 210 nm, with a melting temperature (Tm) of 65.1 ± 0.6 °C, suggestive of parallel G4 (Vorlíčková et al. 2012; Del Villar-Guerra et al. 2017). In NaCl or LiCl, the structure shifted toward a less stable hybrid G4, with Tm values of ∼40°C and ∼45°C, respectively. In contrast 5’-(AAGGG)_8_-3’ CD spectrum displayed a minimum at ∼250 nm, a prominent positive peak near 220 nm, and a shoulder around 230 nm, a pattern characteristic of poly(dA)-rich sequences rather than G4s (Vorlíčková et al. 2012), with a Tm > 95°C under all tested ionic conditions. **B.** *In vitro* transcription assay of CTTTT- or CCCTT-containing vectors showing a repeat size- and motif-dependent reduction of RNA transcription. The transcriptional products were separated on a denaturing gel along with a ssRNA ladder. Full transcripts (arrows) were densitometrically quantified and are represented as single data points, mean and standard error. AU =arbitrary units. Data were analysed using one-way ANOVA followed by Dunnett’s T3 multiple comparisons post-hoc test. n=2 technical replicates. **C.** Schematic representation of Firefly luciferase reporter vector containing increased sizes of non-pathogenic CTTTT or pathogenic CCCTT repeat (left) and quantification of relative Firefly to Renilla luminescence of cells co-transfected with Firefly and Renilla luciferase vectors (right). RLU= relative luminescence units. Data are shown as single data points, mean and standard error. Data were analysed using one-way ANOVA followed by Dunnett’s T3 multiple comparisons post-hoc test. Experiments were performed in triplicate on four independent occasions. **D.** Schematic representation of pGint fluorescence reporter vector containing increased sizes of non-pathogenic CTTTT or pathogenic CCCTT repeat (left) and relative EGFP to mCherry fluorescence quantification of HEK cells co-transfected with pGint and mCherry vectors (right). Data are shown as single data points, mean and standard error. Data were analysed using one-way ANOVA followed by Dunnett’s T3 multiple comparisons post-hoc test. Experiments were performed in triplicates or quadruplicates on three independent occasions.

### RFC1 repeat expansions impair transcription and gene expression in a size- and motif- dependent manner *in vitro*

The above data suggested a capacity of CANVAS repeat expansions to disrupt *RFC1* gene expression. To test this, we generated constructs with incrementally larger expansions of either non-pathogenic (CTTTT) or pathogenic (CCCTT) repeat motifs, ranging from 12 to ∼70 units. These were cloned downstream of a T7 promoter, in the same orientation as they are found on the *RFC1* coding strand. Indeed, in an *in vitro* transcription assay, pathogenic CCCTT repeats were transcribed less efficiently than non-pathogenic CTTTT repeats of similar sizes (**Fig. 1B**). The decrease in transcription was larger for longer repeats, suggesting that large CCCTT repeat expansions might affect RNA polymerase efficiency and processivity.

The transcriptional effect of pathogenic CCCTT expansions was further confirmed using gene reporter vectors overexpressed in HEK293 cells. In a Firefly Luciferase system, CCCTT but not CTTTT expansion led to a size-dependent reduction of normalised expression of a downstream reporter (**Fig. 1C**). Similarly, in a two-exon fluorescent reporter system (pGint), normalised EGFP expression was significantly reduced by the presence of 48 or more intronic CCCTT, but not CTTTT, repeats (**Fig. 1D**).

These results suggest that the pathogenic CCCTT repeat can lead to reduced transcription and gene expression *in vitro*. While technical reasons prevent us from generating constructs with pathogenic expansion sizes in the range observed in CANVAS patients (i.e., greater than ∼250 repeats), we note that the transcriptional effects in these *in vitro* systems are already detectable for much smaller expansions.

### *RFC1* repeat expansions are associated with decreased *RFC1* expression in the cerebellum

To confirm the relevance of our *in vitro* findings in CANVAS patients, we examined the association between AAGGG (CCCTT) expansion and *RFC1* expression in post-mortem brain tissue from biallelic and monoallelic *RFC1* expansion carriers.

First, we performed bulk RNA-seq on cerebellar hemispheres and frontal cortex from CANVAS patients (N=5 and N=6, respectively) and healthy controls (N=4 and N=5, respectively). Clinical and pathological details are summarised in **Supplementary Table 1**.

Principal component decomposition revealed that RNA integrity number (RIN) had a major effect on sample variability, and was highly correlated with PC1, while there was limited clustering in terms of disease status (**Supplementary Fig. 2A**). Using standard package DESeq2, we implemented a model that included RIN and sex as covariates. Based on a false-discovery rate (FDR) of 5%, 28 genes were upregulated and four were downregulated in CANVAS cerebellar hemispheres (**Fig. 2A and Supplementary Table 2**), while four genes were upregulated and one gene was downregulated in frontal cortex tissue (**Supplementary Fig. 2B and Supplementary Table 2**). For upregulated genes in cerebellar hemispheres, Gene Ontology (GO) analysis showed an enrichment of terms related to innate immune responses and microglial activation (**Fig. 2B**). These data are consistent with a recent transcriptomic study that identified activation of immune and inflammatory pathways across different forms of genetic ataxia (Chen et al. 2025).

**Figure 2.**
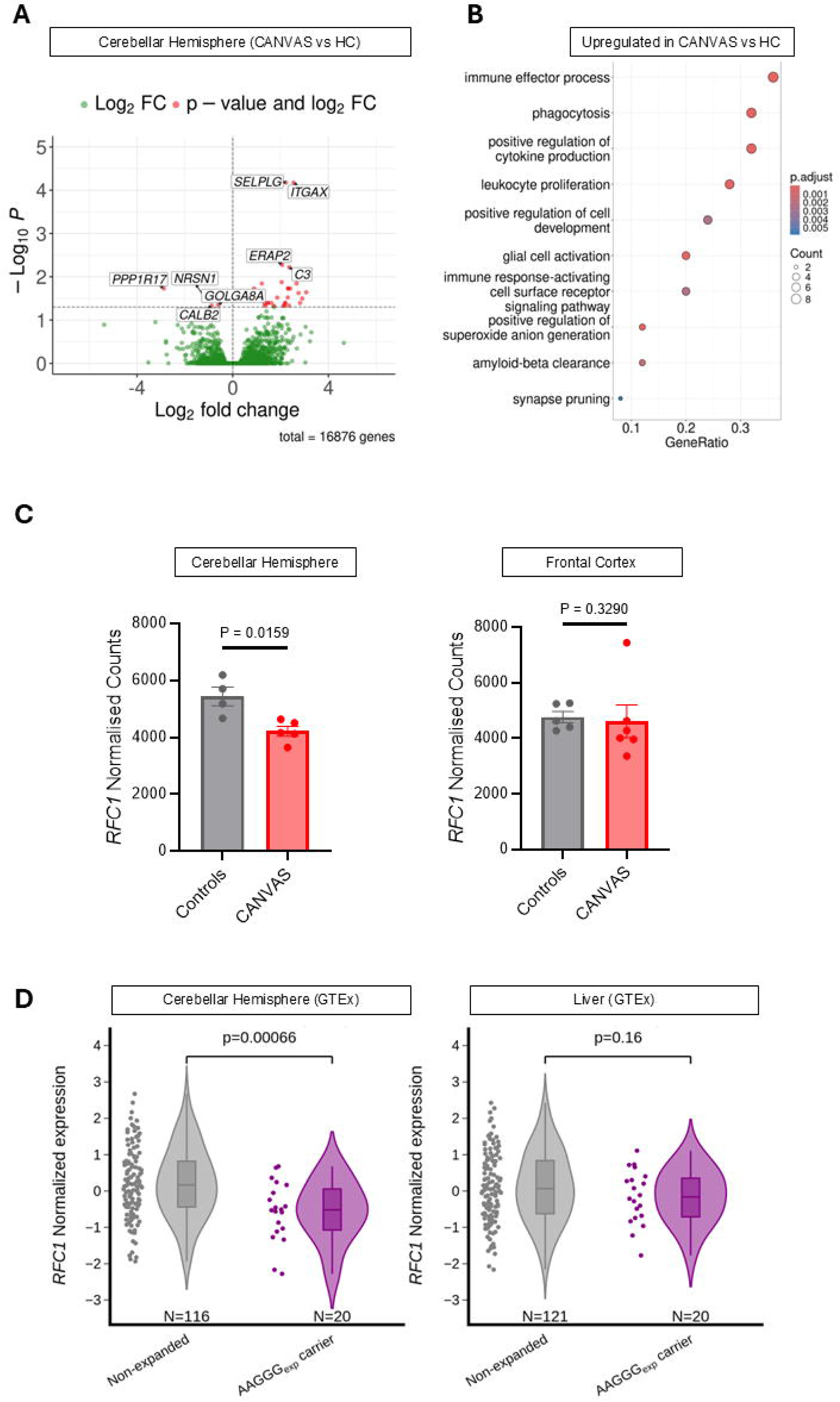
*RFC1* repeat expansions are associated with decreased *RFC1* expression in the cerebellum. **A.** Volcano plot showing differentially expressed (28 upregulated and 4 downregulated) genes in CANVAS (n=5) vs healthy controls (n=4) cerebellar hemispheres. **B.** Gene Ontology (GO) analysis of upregulated genes in CANVAS cerebellar hemispheres showing an enrichment of terms related innate immune responses and microglial activation. The ten most significant GO pathways are shown. **C.** Normalised read counts of *RFC1* in CANVAS vs control cerebellar hemispheres and frontal cortex showing reduced expression of RFC1 in cerebellum but not frontal cortex in CANVAS samples. Data are shown as single data points, mean and standard error. The statistical significance was calculated using unpaired t test analysis. **D.** Violin plots of *RFC1* normalised expression (see methods) in healthy controls part of the GTEx project stratified according to the AAGGG expansion carrier status. The presence of AAGGG pathogenic repeat is associated with reduced expression of *RFC1* in cerebellum but not in liver tissue. The statistical significance was calculated using unpaired t test analysis. HC: healthy controls; FC: fold change.

Expression of *RFC1* was not significantly changed at FDR of 5%. However, when looking specifically at RIN-corrected expression of different subunits of the RFC complex, *RFC1* showed a 25% reduction in cerebellum, but not in frontal cortex, in CANVAS patients vs controls (**Fig. 2C**), while expression levels of the other four subunits (*RFC2-5*) were unchanged (**Supplementary Fig. 2C**). Exon usage of *RFC1* was similar in CANVAS patient tissue vs controls (**Supplementary Fig. 2D**).

While interesting, we interpreted these results with caution due to the limited number of postmortem brains analysed and the generally low RIN (range = 2-5), which is particularly relevant to the quantification of longer transcripts, such as *RFC1*.

Therefore, we turned to large publicly available transcriptomic datasets of high quality (RIN > 5) brain samples to assess the association between the presence of heterozygous pathogenic *RFC1* expansions and *RFC1* expression in affected (cerebellum) and unaffected tissues. Out of 264 individuals on GTEx (v.10) with available RNA-seq and short-read whole genome sequencing data for cerebellum, 20 (7.6%) were found to carry one AAGGG (pathogenic) repeat expansion in heterozygous state with either a non-expanded allele or a non-AAGGG (non-pathogenic) expansion, 128 (48.5%) carried at least one non-AAGGG (non-pathogenic) repeat expansion, while 116 (43.9%) had two non-expanded alleles. Compared to non-expanded individuals, AAGGG expansion carriers have lower expression of *RFC1* in cerebellum (P=0.0007) but not in the liver (**Fig. 2D**) or in other tissues (**Supplementary Table 3**).

To gather further insight on the tissue-specificity of the association between AAGGG expansion and *RFC1* expression, we expanded our analysis to the entire GTEx dataset, including samples without short-read WGS data. Leveraging all available SNPs within the *RFC1* gene, we identified a region extending between SNPs rs13147094 and rs13142220 (chr4:39309262-3935381), corresponding to a linkage disequilibrium block and containing the SNP rs6815219, here used as a proxy for the whole haplotype (**Supplementary Fig. 3A**). Next, by leveraging long read sequencing data from the UK 100K Genome Project we showed that the linkage region acts as a permissive haplotype associated with expansion of different repeat motifs, including the pathogenic AAGGG repeat (**Supplementary Fig. 3B**). Interestingly this expansion-permissive haplotype, including the tagging SNP rs6815219, acts as an expression quantitative trait loci (eQTL) associated with reduced expression of *RFC1* (**Supplementary Fig. 3C**). Notably, the effect is much stronger in cerebellum (N=264), which is primarily affected in this condition, compared to any other tissue available on GTEx and the magnitude is larger for carriers of pathogenic AAGGG vs. non-pathogenic AAAAG expansion (**Supplementary Fig. 3D**). Unfortunately, tissue from dorsal root ganglia or vestibular ganglia were not available on GTEx, so we could not test whether a similar effect could be observed in these structures typically affected in CANVAS.

The association between *RFC1* expression and the repeat-associated haplotype was replicated in the Metabrain dataset (de Klein et al. 2023), which is a meta-analysis of gene expression studies. This dataset includes cerebellum samples from GTEx (N=118), AMP-AD MAYO (N=256), TargetALS (N=72), and UCLA_ASD (N=46), totalling 492 samples (P=3.5 ×10¹). The association was also confirmed when analysing the AMP-AD MAYO dataset separately (P=0.0021).

Overall, studies on post-mortem brain tissue suggest that pathogenic *RFC1* expansions are associated with a small but consistent reduction of *RFC1* transcript levels in CANVAS patients and *RFC1* expansion carriers, which is specific, or at least more prominent, in cerebellar tissue.

### *RFC1* repeat expansions lead to decreased *RFC1* expression in iPSC-derived sensory and motor neurons

We next set out to explore whether reduced *RFC1* expression is also detectable in patient-derived cell lines.

Previous studies (Cortese et al. 2019; Gisatulin et al. 2020a; Maltby et al. 2024) showed that *RFC1* transcript levels did not differ between fibroblast or lymphoblastoid cell lines from CANVAS patients and controls. Since sensory neuronopathy is a consistent feature of RFC1 CANVAS, and it is possible that sensory neurons are more susceptible to changes in *RFC1* expression compared to other cell types, we optimised an inducible differentiation protocol to generate highly pure sensory neuronal cultures (Nickolls et al. 2020). Moreover, we generated isogenic controls from two CANVAS patient iPSC lines by deleting the AAGGG repeat and its flanking AluSx3 element (AluSx3_exp_) (**Fig. 3A**). We were able to produce both heterozygous AluSx3_exp_ +/- and, for the first time, homozygous AluSx3_exp_ -/- corrected lines, as confirmed by PCR and long read sequencing (**Supplementary Fig. 4A-D**), which together with uncorrected (AluSx3_exp_ +/+) CANVAS patient-derived lines were differentiated into sensory neurons.

**Figure 3.**
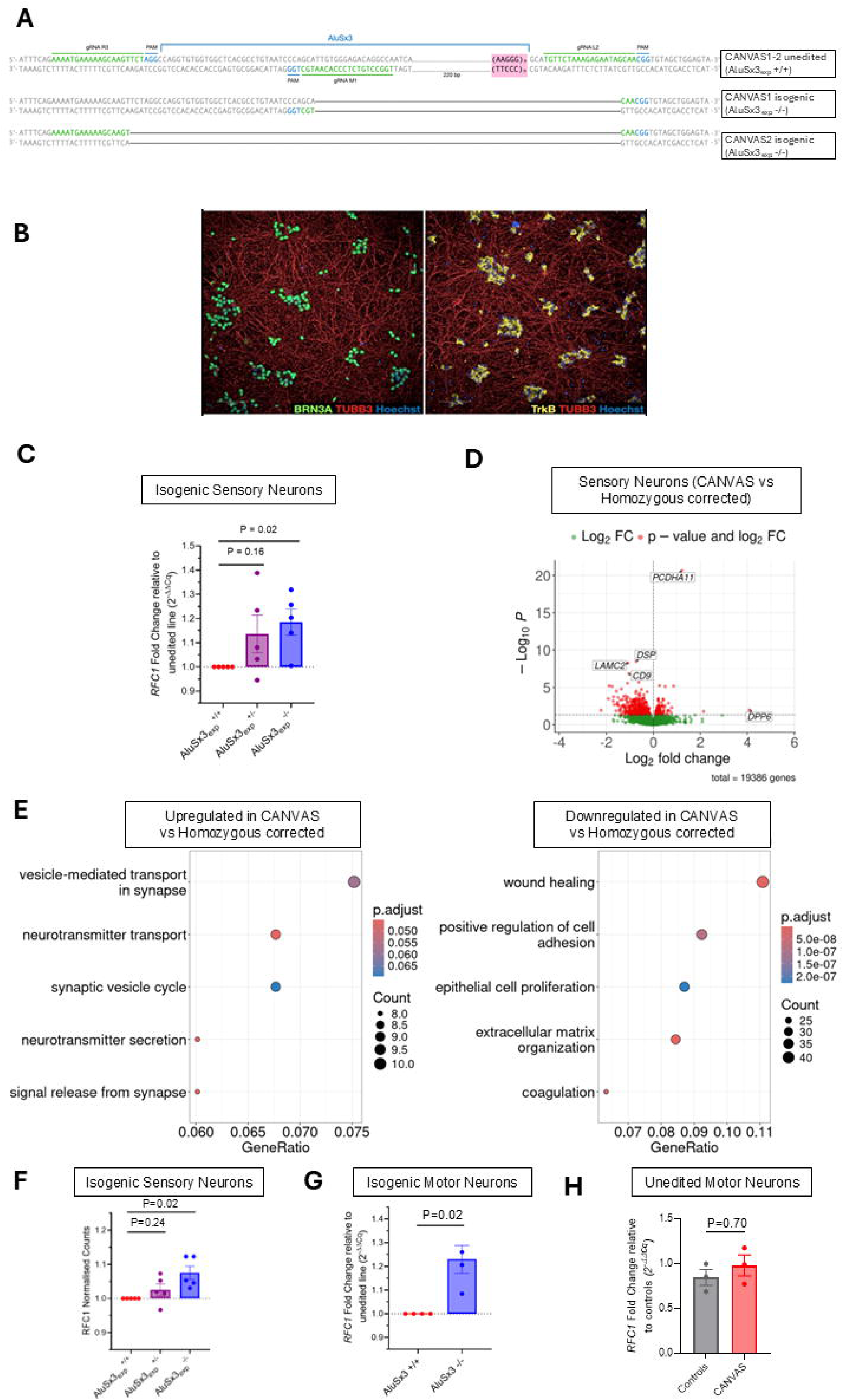
*RFC1* repeat expansions lead to decreased *RFC1* expression in iPSC-derived sensory and motor neurons. **A.** DNA sequence (based on Nanopore long-read sequencing, more details in **Supplementary** Fig. 4C) of CANVAS parent lines (AluSx3_exp_ +/+) and their corresponding biallelic edited isogenic lines (AluSx3_exp_ -/-) confirming successful editing. Target nucleotides for sgRNAs are in green, while PAM (Protospacer Adjacent Motif) sites are in light blue. For both patient lines, CANVAS1 and CANVAS2, the same L2 guide was used as 5’ sgRNA, whereas either M1 or R3 were used as 3’ sgRNA guide. **B.** Representative immunohistochemistry images showing that forced (doxycycline inducible) expression of Ngn2 and Brn3a produces highly pure sensory neuronal cultures that stain positive for β3-Tubulin (TUBB3), Brn3a (POU4F1) and TrkB (NTRK2). Scale bar= 100 μm. **C.** Quantitative RT-PCR for *RFC1* expression in AluSx3_exp_ +/- and AluSx3_exp_ -/- relative to their uncorrected (AluSx3_exp_ +/+) day 7 iPSC-derived sensory neurons. Relative expression levels (RQ) of *RFC1* are normalized to *TBP* using the 2^–ΔΔCt method (Livak & Schmittgen, 2001). Paired analysis of 3 and 2 separate differentiations (batches) for patients CANVAS1 and CANVAS2 lines, respectively. **D.** Volcano plot of differentially expressed genes in CANVAS patients (AluSx3_exp_ +/+) vs biallelic isogenic control (AluSx3_exp_ -/-) iPSC-derived sensory neurons **E.** Gene Ontology (GO) analysis of upregulated (left) and downregulated (left) genes in CANVAS patients (AluSx3_exp_ +/+) vs biallelic isogenic control (AluSx3_exp_ -/-) iPSC-derived sensory neurons. The five most significant GO pathways are shown. **F.** Ratio of Normalised read counts for RFC1 mRNA in heterozygous (AluSx3_exp_ +/-) and homozygous (AluSx3exp -/-) isogenic control iPSC-derived sensory neurons (day 7) compared to their unedited CANVAS (AluSx3_exp_ +/+) parent line. Technical replicates n=3 for CANVAS1 and n=2 for CANVAS2 lines, data are shown single data points, mean and standard error. The statistical significance was calculated using paired t test analysis. **G-H.** Quantitative RT-PCR for *RFC1* expression in iPSC-derived motor neurons from AluSx3-/- relative to their uncorrected (AluSx3_exp_ +/+) (**G**) and in healthy controls vs CANVAS-derived lines (**H**). Relative expression of RFC1 was normalized to TBP and measured using the 2^–ΔΔCt method.

To limit bottleneck-associated variability, uncorrected AluSx3_exp_ +/+ cells were processed in the same way as the isogenic lines, including selection down to single cells followed by clonal expansion. Neurons appeared strongly positive for BRN3A and TrkB, suggestive of a mechanoreceptor identity (**Fig. 3B**). Morphologically, lines were indistinguishable from one another throughout the 7 days of differentiation (**Supplementary Fig. 5A**).

Notably, *RFC1* expression, as measured by qPCR, was 15% lower in CANVAS (AluSx3_exp_+/+) samples when matched to their isogenic (AluSx3_exp_ -/-) controls (P=0.024). *RFC1* expression in heterozygous corrected AluSx3_exp_ +/- samples also appeared to be reduced but the difference was not statistically significant (P=0.16) (**Fig. 3C**).

To better understand the transcriptomic profile of CANVAS sensory neurons, we performed RNA-seq in AluSx3_exp_ +/+, AluSx3_exp_ +/- and AluSx3_exp_ -/- lines. PCA showed clustering of samples primarily according to stages in neuronal differentiation (iPSCs vs. Day 7 Sensory Neurons), but also relative to patient of origin, differentiation batch, and (to a minor degree) AluSx3_exp_ correction status (**Supplementary Fig. 5B**). The sensory neurons thus had transcriptomic hallmarks (PIEZO2+ and NTRK2+) suggestive of human mechanoreceptors (**Supplementary Fig. 5C**) (Nickolls et al. 2020). Differential gene expression analysis between CANVAS (AluSx3_exp_ +/+) and isogenic control (AluSx3_exp_ -/-) neurons identified 151 upregulated and 406 downregulated genes in CANVAS (**Fig. 3D** **and Supplementary Table 4**). For upregulated genes, GO terms were enriched for neurotransmitter transport and synaptic processes, whereas for downregulated genes there was enrichment for terms associated with cell adhesion (**Fig. 3E**), adding further evidence that synaptic dysfunction and reduced cellular maintenance could be downstream mechanisms potentially contributing to neurodegeneration in CANVAS (Maltby et al. 2024). Similar findings were obtained when comparing AluSx3_exp_ +/+ with both heterozygous and homozygous corrected lines together (AluSx3_exp_ +/- and -/-) (**Supplementary Fig. 6A and Supplementary Table 4**).

Importantly, *RFC1* expression in AluSx3_exp_ +/+ lines was confirmed to be reduced relative to matched AluSx3_exp_ -/- corrected lines (P<0.01) but not to AluSx3_exp_ +/- lines (**Fig. 3F**), following FDR correction. Interestingly, when looking specifically at expression of other RFC subunits, AluSx3_exp_ +/+ lines were found to have lower expression of all five subunits, although apart from RFC1 this was only statistically significant for RFC3 and RFC5 (**Supplementary Fig. 6B**). These results suggest a possible shared regulation of the transcription of related genes of the replication factor complex in sensory neurons.

Since involvement of lower motor neurons has been reported in CANVAS (Huin et al. 2021), we also differentiated CANVAS (AluSx3_exp_ +/+) and isogenic (AluSx3_exp_ -/- ) iPSC lines, along with three non-isogenic controls, into spinal lower motor neurons using dual-SMAD inhibitors (Pagliari et al. 2025). No differences were observed at any stage of the differentiation process between CANVAS and control lines in terms of morphology and neurite outgrowth (**Supplementary Fig. 7A**). Around 70% of cells expressed the lower motor neuron-specific marker HB9 (**Supplementary Fig. 7B**).

Similarly to sensory neurons, *RFC1* expression in iPSC-derived lower motor neurons appeared ∼23% lower in CANVAS (AluSx3_exp_ +/+) (P=0.025), as measured by qPCR (**Fig. 3G**). Notably a reduction of *RFC1* expression could only be detected when comparing CANVAS lines with their isogenic corrected counterpart, but not when comparing CANVAS lines with non-isogenic controls (**Fig. 3H**), highlighting the critical utility of isogenic controls in eliminating confounding genetic background effects.

Finally, *RFC1* transcript processing – assessed through RNA-seq in sensory neurons (**Supplementary Fig. 8A**) and amplicon based targeted long read sequencing in sensory and motor neurons (**Supplementary Fig. 8B-C**) – appeared unchanged in CANVAS vs isogenic and non-isogenic controls.

Overall, results from iPSC-derived sensory and motor neurons confirm our observations in post-mortem brains and *in vitro* transcription models that pathogenic AAGGG repeat expansions are associated with reduced *RFC1* expression, and conversely, that biallelic excision of the pathogenic repeat and flanking Alu element is sufficient to increase *RFC1* expression in CANVAS lines by ∼20%.

### CANVAS cell lines show increased sensitivity to platinum-induced DNA damage, supporting an impairment of RFC1 DNA repair function

We next set out to investigate whether *RFC1* repeat expansions can affect RFC1 function using patient-derived cell lines.

Notably, 80% *RFC1* knockdown through *RFC1*-targeting shRNA in a wild-type iPSC sensory neuron line reduced neuronal survival compared to a scrambled shRNA, as monitored by continued live-cell imaging up to 21 days (**Supplementary Fig. 9A-B**). However, the survival of sensory neurons harbouring CANVAS-associated repeats was similar compared to isogenic control lines (data not shown). Indeed, the magnitude of the reduction of *RFC1* in CANVAS neurons is small, and it was unclear whether it would be significant enough to produce a cellular phenotype during the 1-month observation period.

Neurons are particularly sensitive to DNA damage and oxidative stress due to their high metabolic demand and transcriptional activity (Welch and Tsai 2022; Caldecott et al. 2022). Therefore, we speculated that the use of an exogenous stressor may be required to reproduce the characteristic cumulative insults of late-onset neurodegenerative diseases and to unmask potential functional differences between CANVAS and control lines. To this regard, we decided to focus on platinum-based drugs because of **1)** the essential role played by RFC-PCNA in the repair of platinum-induced DNA damage through nucleotide excision repair (Araújo et al. 2000), and **2)** the known sensitivity of sensory neurons to this type of DNA damaging agent (Staff et al. 2019)

iPSC-derived neurons from CANVAS (AluSx3_exp_ +/+), isogenic controls (AluSx3_exp_ -/-), and non-isogenic controls, were therefore treated with increasing doses of cisplatin. An increased rate of apoptosis, defined by the percentage of cleaved Caspase-3 positive nuclei, was observed when comparing motor neurons from two CANVAS patients vs three non-isogenic controls after 24 h treatment with 5 and 10 µM cisplatin (**Fig. 4A-B**). Notably, biallelic excision of the AluSx3_exp_ was able to rescue this phenotype in CANVAS lines, leading to a marked reduction of the number of apoptotic cells in cisplatin-treated AluSx3_exp_ -/- iPSC-derived motor neurons compared to their unedited AluSx3_exp_ +/+ parent lines (**Fig. 4C**). Cisplatin treated isogenic AluSx3_exp_ +/- and -/- iPSC sensory neurons also showed a consistent trend towards a reduction of the apoptotic rate compared to unedited AluSx3_exp_ +/+ CANVAS parent lines, reaching significance after 24 h of 5 µM cisplatin treatment (**Fig. 4D**).

**Figure 4.**
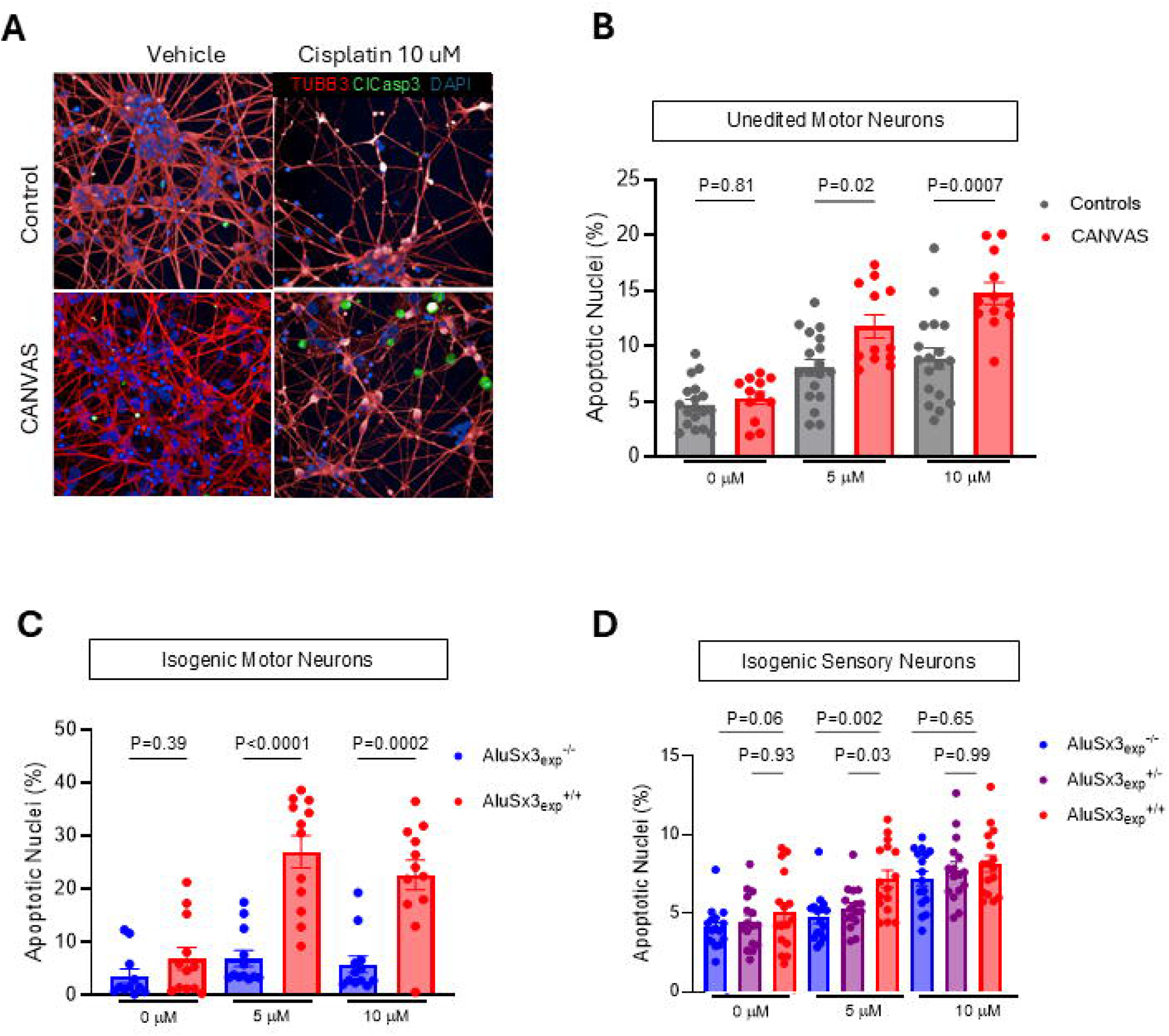
CANVAS iPSC-neurons show increased sensitivity to platinum-induced DNA damage, supporting an impairment of RFC1 DNA repair function. A. Representative images of immunostaining on iPSC-derived motor neurons showing an increased number of cleaved Caspase-3 positive (green) nuclei in CANVAS patients compared to controls after cisplatin treatment. Scale bar = 50 μM **B-D.** Increased apoptosis rate in iPSC-derived motor (**B-C**) and sensory (**D**) neurons from CANVAS patients compared to isogenic and non-isogenic controls after 24h treatment with cisplatin 5 mM or 10 mM. The apoptotic rate was calculated as the number of nuclei positive for cleaved Caspase-3 out of the total number of neuronal nuclei. Data are shown as single data points, mean and standard error. The statistical significance was calculated using Brown-Forsythe and Welch ANOVA test followed by Dunnett’s T3 multiple comparisons post-hoc test. All experiments were replicated twice.

To further expand our results on more scalable lines and gain further insight into the possible dysfunction of DNA damage response, we leveraged lymphoblastoid cell lines (LCLs) from five CANVAS patients and five healthy controls. CANVAS LCLs exhibited reduced viability compared to control lines upon treatment with increasing concentrations of cisplatin (**Fig. 5A**). Activation of the DNA damage response following cisplatin treatment was detected in both CANVAS and control LCLs, as indicated by increased expression of p53, without significant difference between patient and control lines. However, this effect was accompanied by earlier activation of apoptosis in patient-derived LCLs after either 24 or 48 h of continuous treatment with cisplatin as showed by an increased level of cleaved Caspase-3 (**Fig. 5B**). While basal levels of the DNA damage marker γH2A.X varied considerably among LCLs, cisplatin treatment further increased γH2A.X levels, with a more pronounced effect observed in CANVAS LCLs (**Fig. 5B**). Similar results were obtained when LCLs were treated with oxaliplatin (**Supplementary Fig. 10A-B**), a related platinum-based compound that is widely used in clinic for the treatment of colorectal cancers, and which can cause neuropathy.

**Figure 5.**
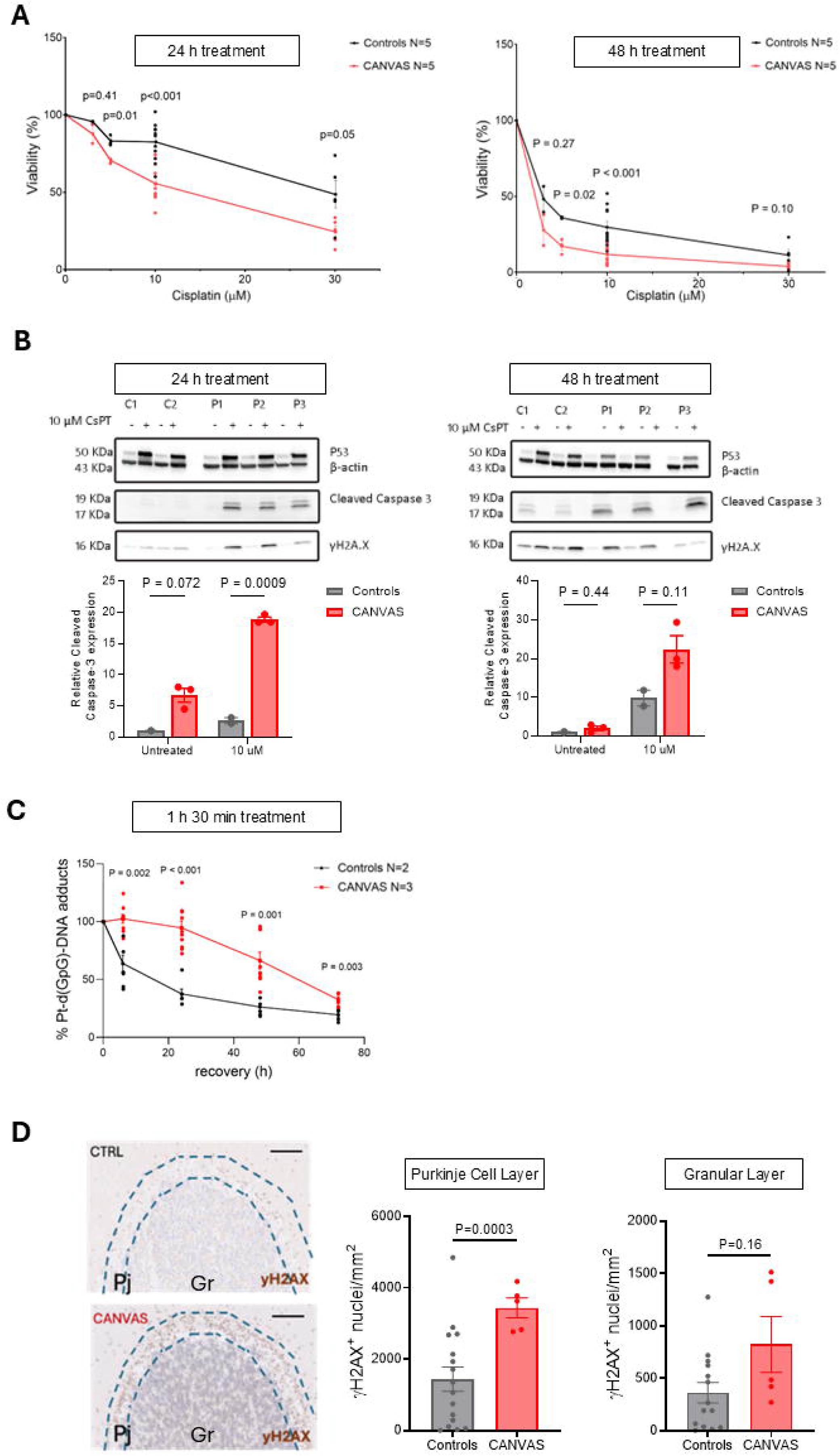
Evidence from CANVAS lymphoblastoid cell lines and post-mortem brain tissue supporting an impairment of RFC1 DNA repair function. A. Viability in CANVAS lymphoblastoid lines (LCLs) is significantly reduced after 24 h (left) and 48 h (right) treatment with increasing doses of cisplatin (from 0 to 30 µM). Data are shown as single data points, mean and standard error. Unpaired t-test with Welch correction was performed to compare two groups assuming unequal variance between the two. Experiments were carried out on 5 CANVAS and 5 control LCLs and each condition was replicated at least three times. **B.** Western blot analysis showing the expression of p53, cleaved Caspase-3 and γH2A.X in LCLs from healthy controls (C1-C2) and CANVAS patients (P1-P2-P3) after 24hr (left) or 48 hr (right) treatment with cisplatin 10 μM. Data are shown as single data points, mean and standard error. Statistical significance was assessed using Brown-Forsythe and Welch ANOVA test followed by Dunnett’s T3 multiple comparisons post-hoc test. **C.** Dot blot quantification of Pt-d(GpG)-DNA adducts in total DNA extracted from CANVAS (red) and healthy controls (black) LCLs upon 1.5h cisplatin pulse administration. The number of LCLs is reported on the right. Individual data points, mean values ± SEM of three different experiments (n=3) are shown. Unpaired t-test with Welch correction was performed to compare two groups assuming unequal variance between the two. **D.** On the left, representative immunohistochemistry image showing increased γH2A.X signal in the post-mortem cerebellum of CANVAS patients compared to healthy controls. Signal intensity was highest at the interface between the molecular and granular layers (highlighted). On the right, quantification of the γH2A.X cell density at the Purkinje cells layer and the granular cells layer, expressed as number of γH2A.X positive nuclei per mm^2^. Data are shown as single data points, mean and standard error. Unpaired t-test with Welch correction was performed to assess the statistical significance. Scale bar: 250 μm

The increased susceptibility of CANVAS cell line to continuous cisplatin treatment was also further confirmed when LCLs were exposed to 1.5 h cisplatin pulse treatment, with CANVAS cells showing reduced viability compared to controls. Additionally, CANVAS LCLs showed marked accumulation in pre-G1 stage at both 24 h and 48 h of recovery (**Supplementary Fig. 10C-D**), consistent with induction of apoptosis. This is confirmed by Annexin V/PI staining showing apoptotic cell accumulation in CANVAS LCLs but not in controls (**Supplementary Fig. 10E**) and increased cleaved Caspase-3 level on Western Blotting (**Supplementary Fig. 10F**).

After establishing an increased sensitivity of CANVAS LCLs to cisplatin and oxaliplatin, we asked whether this was due to an impaired repair of platin-induced DNA damage. In the cell, the platinum moiety forms covalent bonds with the N7 position of purine bases, predominantly generating 1,2-intrastrand crosslinks (65% GpG and 25% ApG), which are also termed platinum-(GpG) DNA adducts (Lemaire et al. 1991). After 1.5 h of cisplatin pulse treatment, while control LCLs efficiently removed DNA damage already at the earliest time points, repair in CANVAS LCLs was significantly slower, reaching levels comparable to control cells only after 72 h (**Fig. 5C**). These data reveal slower kinetics of DNA repair in patient-derived LCLs, leading to accumulation of DNA damage and activation of apoptosis, supporting the presence of RFC1 dysfunction and an impaired DNA damage response in CANVAS.

We then investigated whether the DNA damage marker yH2A.X also accumulates in post-mortem brains from CANVAS patients. Cerebellum formalin fixed paraffin embedded sections from post-mortem brains were available from six CANVAS patients and 16 age-matched controls (**Supplementary Table 1**). While the nearly complete loss of Purkinje cells in CANVAS patients prevents analysis of γH2A.X levels in this specific cell type, we observed an increased γH2A.X signal in the Purkinje cell layer, primarily affecting the residual supporting Bergmann glia, but not the granular cell layer in CANVAS cases compared to controls (**Fig. 5D**).

In conclusion, the increased sensitivity of CANVAS cells to DNA-damaging agents and the slower repair of DNA damage, together with the accumulation of DNA damage markers in post-mortem cerebellar tissue, supports a role for reduced RFC1 function and defective DNA repair in CANVAS pathogenesis, which can be rescued in cellular models by biallelic excision of the expanded repeat.

### Neuronal knockdown of the *Drosophila RFC1* ortholog *Gnf1* causes late-onset motor defects and DNA damage accumulation

To explore the consequences of partial loss of RFC1 function *in vivo*, we developed a *Drosophila melanogaster* model by performing shRNA-mediated knockdown of *Gnf1*, the fruit fly ortholog of *RFC1* (**Supplementary Fig. 11A**) (Mossi et al. 1997).

Consistent with the lethality of homozygous *Gnf1* null alleles and the essential roles of Gnf1 in DNA replication and cell cycle progression (Tsuchiya et al., 2007), global knockdown of *Gnf1* led to complete lethality. We then generated a neuron-specific knockdown model by using the *nsyb*-Gal4 driver to express *Gnf1* shRNA specifically in post-mitotic neurons, resulting in a > 50% reduction in *Gnf1* mRNA expression in dissected adult brain tissue (**Supplementary Fig. 11B**).

Recent single-cell sequencing approaches have revealed widespread expression of *Gnf1* in *Drosophila* neurons (**Supplementary Fig. 11C**) (Davie et al. 2018). To determine whether pan-neuronal *Gnf1* knockdown yielded motor phenotypes relevant to CANVAS, we assayed locomotor activity in male flies at various time-points using the *Drosophila* Activity Monitor (DAM) system (**Supplementary Fig. 11D**) (Pfeiffenberger et al. 2010). We observed no significant change in the locomotor activity of 3-7 and 21-23 day old *Gnf1* knockdown flies (**Fig. 6A- B,** and **Supplementary Fig. 11E-F**). However, consistent with the late-onset of ataxia in CANVAS patients (mean = 54 years; IQR = 49-61) (Currò et al. 2023), in aged (40- 42 day old) flies, pan-neuronal *Gnf1* knockdown induced a significant reduction in total locomotor activity compared to driver- and transgene-alone controls (**Fig. 6C**, and **Supplementary Fig. 11G**). We also found that pan-neuronal *Gnf1* knockdown significantly shortened the lifespan of adult flies (P=0.001), with death of *Gnf1* knockdown flies first observed at ∼ 20 days compared to ∼ 40 days in controls (**Fig. 6D**). Hence, pan-neuronal *Gnf1* knockdown causes late-onset motor and organismal defects, similar to those observed in CANVAS patients.

**Figure 6.**
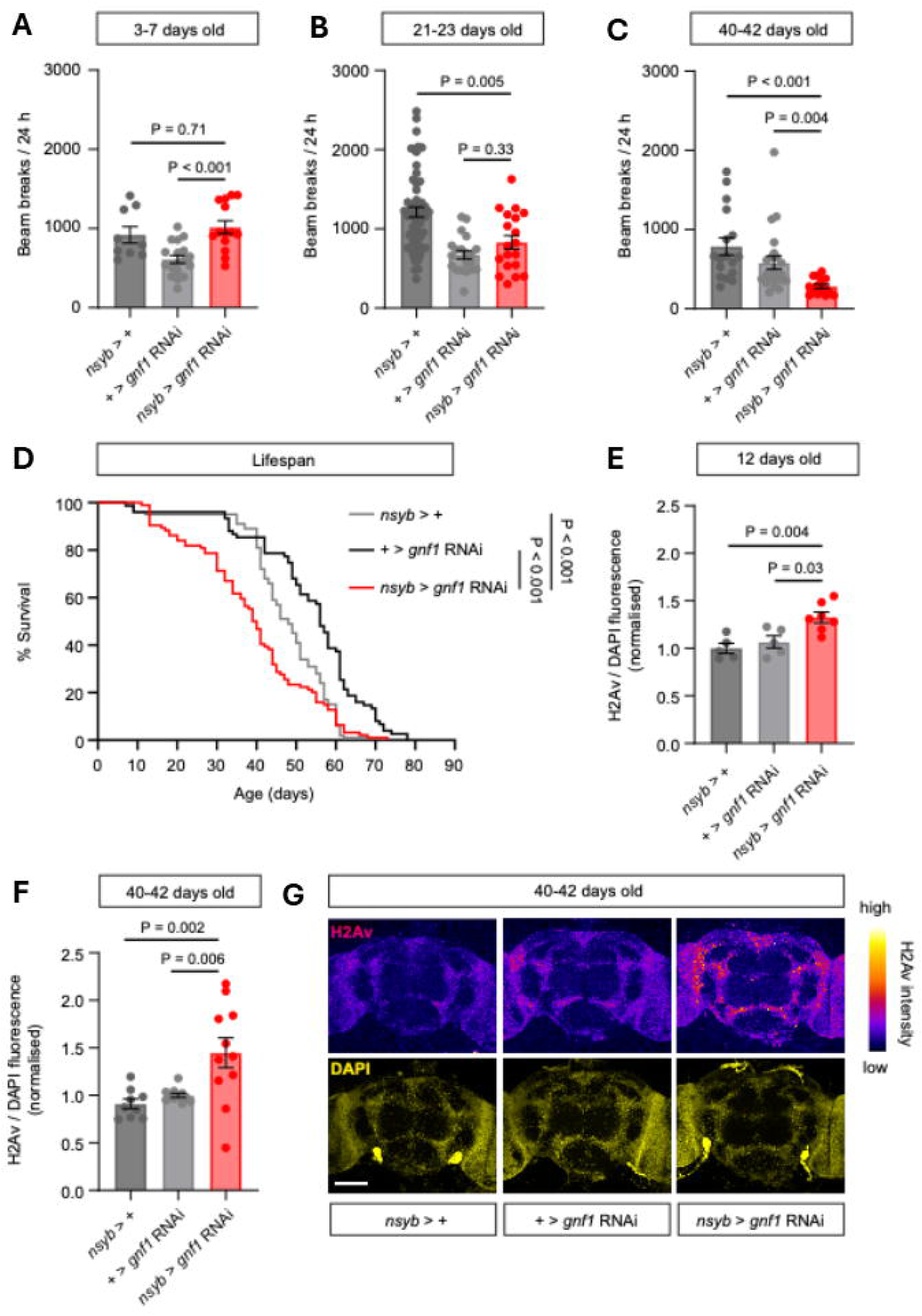
Neuronal knock-down of the *Drosophila* RFC1 ortholog Gnf1 causes late-onset motor defects and DNA damage accumulation. **A-C.** Locomotor activity in adult male driver/shRNA alone controls flies, or flies expressing Gnf1 shRNA in post-mitotic neurons (*nsyb* > *gnf1* shRNA), of varying ages, quantified as beam breaks across a 24 h period in the *Drosophila* activity monitor (DAM). **A**: 3-7 days old flies (n=9-18); **b**: 21-23 days old flies (n=19-60); and **C**. 40-42 days old flies (n=15-23). **D.** Survival curve showing the percentage survival of *nsyb* > *gnf1* shRNA flies compared to controls during normal aging. n=100 per genotype. **E-G**. DNA damage accumulation in control and neuronal Gnf1 knockdown adult male fly brains, quantified as the ratio of H2Av immuno-fluorescence to DAPI fluorescence at 12 days old (**e**: n=5-7) and 40-42 days old (**f**: n=9-11). **G**. Representative confocal images illustrating H2Av and DAPI staining in 40-42 days control and neuronal *Gnf1* knockdown adult male fly brains. H2Av immuno-fluorescence is color-coded to indicate signal intensity. Scale: 100 µm. Error bars in **A-C**, **E**, **F**: standard error of the mean (SEM). Central line in dot plots: mean. P values are indicated, acquired via Brown-Forsythe and Welch ANOVA test with Dunnett’s T3 multiple comparisons post-hoc test (A,E,F) or Kruskal-Wallis test with Dunn’s T3 multiple comparisons post-hoc test (B, C) or Mantel-Cox Log rank test (D)

To assess the relationship between Gnf1 and DNA damage in *Drosophila* neurons, we immuno-stained brains of neuron-specific *Gnf1* knockdown flies and controls for H2Av, which, similarly to γH2A.X in humans, acts as a *Drosophila*-specific marker for DNA damage (Lake et al. 2013). We selected two adult ages at which to perform these experiments: 12 days old (when no mortality or locomotor defects are observed in *Gnf1* knockdown flies; **Fig. 6A,D**), and 40-42 days old (when locomotor defects and substantial mortality are apparent in *Gnf1* knockdown flies; **Fig. 6C-D**). Interestingly, we observed a significant increase in neuronal H2Av staining in *Gnf1* knockdown flies compared to controls at both ages (**Fig. 6E-G**). DNA damage appeared higher in aged compared to young *Gnf1* knockdown flies, as a proportion of 40-42 day old neuron-specific *Gnf1* knockdown flies exhibited > 2-fold increases in neuronal H2Av levels relative to controls, while increases of this magnitude were not observed in corresponding younger flies (**Fig. 6E-G**).

Given that our human iPSC-derived CANVAS models consistently demonstrated increased sensitivity to the DNA damaging agent cisplatin, we also examined the effects of cisplatin in neuron-specific *Gnf1* knockdown flies. Ingestion of cisplatin robustly reduced lifespan in control flies, with all flies dying prior to the age of 33 days old (**Supplementary Fig. 12A**; compare with **Fig. 6D**). Importantly, the detrimental effect of cisplatin was significantly enhanced by simultaneous neuronal *Gnf1* knockdown (**Supplementary Fig. 12A**). We also observed a novel locomotor phenotype that emerged prior to early mortality in 12 day old neuronal *Gnf1* KD flies fed cisplatin (**Supplementary Fig. 12B**), characterised by a robust advance of evening anticipation – a circadian clock-driven increase in locomotion prior to lights-off (Grima et al. 2004) – and reduced startle responses to lights-off (ZT12-14) (**Supplementary Fig. 12C, D**). These findings suggest that inducing DNA damage when *Gnf1* expression is reduced may further perturb the activity of pre-motor and sensory pathways that temporally pattern locomotor activity in *Drosophila*.

Together, these data suggest that neuronal-specific knockdown of the *RFC1* ortholog in flies leads to accumulation of neuronal DNA damage, late-onset progressive motor impairment and reduced survival, which are exacerbated by treatment with DNA damaging cisplatin.

### Heterozygous AAGGG *RFC1* repeat expansions increase the susceptibility to oxaliplatin-induced peripheral neuropathy in cancer patients

Platinum-based compounds are important chemotherapeutic agents currently used in the treatment of various malignancies. Despite their fundamental impact in increasing survival of cancer patients, chemotherapy-induced peripheral neuropathy (CIPN) is a common and dose-limiting adverse effect of both cisplatin and oxaliplatin (Bonomo and Cavaletti 2021).

Platinum-induced neuropathy is primarily a sensory neuronopathy, and as such it presents with similar clinical manifestations to CANVAS (Staff et al. 2019). Obviously, we could not directly assess whether the detrimental effect of platinum compounds observed in CANVAS cell lines and *Drosophila* also occurs in CANVAS patients by giving them platinum drugs. However, under a theoretical loss-of-function model, it would be expected that each pathogenic expanded allele leads to reduced *RFC1* expression and function, and therefore that heterozygous AAGGG expansion carriers may be more susceptible to CIPN compared to non-carriers. To test this hypothesis, we leveraged an international cohort of 361 colorectal cancer patients treated with oxaliplatin and assessed the association between *RFC1* expansion and presence and severity of the CIPN.

We selected only patients without a previous history of neuropathy, no prior exposure to chemotherapy, and who had received a cumulative dose of at least 1,000 mg of oxaliplatin. All selected patients were clinically graded by a trained neurologist using the validated clinical version of the Total Neuropathy Score scale (TNSc) and genetically tested for the presence of one AAGGG expanded allele. Importantly, clinical teams were blind to patient *RFC1* genotypes at the time of neurological assessment. We identified 34 heterozygous AAGGG expansion carriers (9.4%), in line with the expected allele frequency in the population, and no individuals carrying biallelic AAGGG expansions. Whereas 82% (269/327) of non-carriers developed any degree of CIPN (TNSc > 0), 100% (34/34) carriers of AAGGG alleles were found to have some degree of clinically detectable neuropathy (P=0.003) (**Table 1**). Moreover, clinically relevant neuropathy (TNSc > 7) was identified significantly more frequently in AAGGG expansion carriers relative to non-carriers (70.6% vs. 45.3%, P=0.006). Using logistic regression to correct for effects of age, sex, and centre, we confirmed that *RFC1* expansions were independently associated with a higher risk of developing clinically relevant neuropathy (OR: 2.86; CI 95%: 1.32-6.21; P=0.008).

**Table 1.**

|  | **Whole Series**<br>(n=361;100%) | **Non-carriers**<br>(n=327;90.6%) | **RFC1 AAGGG**<br>**expansion carriers**<br>(n=34; 9.4%) | **P-value** |
| --- | --- | --- | --- | --- |
| **Age** (mean ± SD) | 64.12 ± 9.65 | 64.02 ± 9.64 | 65.06 ± 9.83 | 0.56 |
| **Sex** (Males) | 64.2% | 64.2% | 64.7% | 1 |
| **TNSc** (median, IQR) | 7 (4-11) | 6 (4-10) | 10 (6-13) | **0.004** |
| **Any Neuropathy**<br>(TNSc>0) | 303 (83.9%) | 269 (82.3%) | 34 (100%) | **0.003** |
| **Clinically Relevant Neuropathy**<br>(TNSc>7) | 172 (47.6%) | 148 (45.3%) | 24 (70.6%) | **0.006** |

TNSc= Total Neuropathy Scale, clinical version

In conclusion, heterozygous AAGGG expansion carriers appear to be at increased risk of developing CIPN when treated with oxaliplatin and, when affected, tend to experience more severe symptoms and signs. These findings provide strong clinical validation for a mechanistic link between *RFC1* expansion and impaired DNA damage response in vulnerable sensory neurons.

## DISCUSSION

Here we demonstrate that CANVAS-causing AAGGG (CCCTT) repeat expansions are associated with tissue- and cell type-specific reductions in the expression of *RFC1* transcripts, along with an impaired RFC1 function and increased sensitivity to DNA damage from platin drugs. These observations in post-mortem brain tissue and patient-derived cell lines were further supported by *in vitro* studies showing the propensity of pathogenic AAGGG repeats to form G-quadruplexes, and the reduced transcription from AAGGG-containing templates, with more pronounced effects at longer repeat lengths and increased G-content of the pathogenic repeat motif.

The initial observation of an apparent lack of reduction of *RFC1* mRNA or protein in replicating cell lines (and one post-mortem brain) has puzzled researchers since the first identification of this novel repeat, challenging the paradigm linking recessive diseases to loss of function of the disease-causing gene. Our study offers a comprehensive explanation for this conundrum. Unlike Friedreich ataxia, where biallelic intronic GAA repeat expansions in *FXN* lead to a widespread and pronounced reduction of FXN (Campuzano et al. 1996; Bidichandani et al. 1998), in CANVAS **1)** expression of *RFC1* appears to be only mildly reduced (∼20%), **2)** the effect of the repeat on gene expression is highly tissue-specific, as it could only be identified in affected cerebellum, but not in other unaffected brain regions or peripheral tissues, and **3)** in cell lines, a significant reduction of *RFC1* transcript was only detected using biallelic isogenic lines differentiated into sensory or motor neurons, which minimised variability in *RFC1* expression arising from replication and differences in the genetic background of non-isogenic lines.

It is perhaps not surprising that a large reduction of *RFC1* is not detectable, as complete RFC1 knockout is incompatible with life, both at the cellular level and in animal models, including mice (https://www.informatics.jax.org/allele/MGI:5495286), *Drosophila* (Tsuchiya et al. 2007), and a recent zebrafish model, where global *RFC1* knockdown led to early lethality along with abnormal cerebellar development (Nobilleau et al. 2025). Importantly, we demonstrate that a ∼50% reduction of *RFC1* in *Drosophila* neurons is sufficient to reproduce the late-onset motor phenotype observed in CANVAS patients, which was accompanied by progressive DNA damage accumulation in the brain.

The basis for the marked tissue specificity of RFC1 downregulation in CANVAS remains unclear. Possible explanations include **1)** an interaction between the *RFC1* expansion and cerebellum-specific transcription factors, enhancers or chromatin accessibility pattern – potentially also common to sensory and vestibular neurons – an interpretation further supported by the observation that the repeat associated haplotype acts as a cerebellum-specific eQTL for *RFC1* expression, **2)** the intense transcriptional, metabolic and electrical activity of these highly active cells, leading to the accumulation of DNA damage and creating a demand for RFC1 that exceeds the capacity of the transcriptional machinery to supply it, and **3)** a possible somatic expansion of the AAGGG repeat in this specific cell type, as showed in affected medium spiny neurons in Huntington’s disease (Handsaker et al. 2025), further decreasing *RFC1* expression in single cells.

The most important implication of our study is for the development of therapies for CANVAS patients. Indeed, biallelic excision of the repeat was sufficient to increase *RFC1* expression by ∼20%, which matches the reduction observed in patient cerebellum. To this regard it is worth noting that, up to this time, only heterozygous corrected lines could be generated, leading to the hypothesis that the AluSx3 element and/or the flanking tandem repeat play an essential role in *RFC1* expression and for cell survival (Davies et al. 2023; Maltby et al. 2024). However, here we show that complete excision of both repeat and AluSx3 element is tolerated and appears safe, at least at the cellular level.

While we cannot currently rule out that additional mechanisms may contribute to CANVAS pathology, including repeat-derived peptides, RNA foci, or broader transcriptional changes impacting neuronal function (Wada et al. 2024; Maltby et al. 2024), our data suggest that therapies aimed at increasing *RFC1* expression/function may represent reasonable approaches, warranting further investigation.

Finally, our study has implications for other common diseases, particularly CIPN. Although carriers of the AAGGG expansion do not typically develop neuropathy or ataxia within their natural lifespan, our data demonstrate that they have a significantly increased risk of developing moderate-to-severe neuropathy when treated with oxaliplatin for an underlying malignancy. Importantly, since oxaliplatin does not effectively cross the blood–brain barrier, it is unlikely to exert comparable toxic effects on Purkinje cells in the cerebellum. CIPN remains the leading cause of discontinuation of life-saving platinum-based chemotherapy in patients with high-prevalence cancers, such as colorectal cancer. While our findings suggest a potential clinical utility for *RFC1* genotyping prior to the initiation of platinum-based chemotherapy to stratify neuropathy risk and possibly guide dose modification or treatment scheduling, independent validation in larger cohorts is required before clinical implementation.

As a first-in-kind study, several limitations should be acknowledged. Due to the modest reduction of *RFC1* mRNA observed in both post-mortem brain tissue and CANVAS cell lines, combined with the suboptimal performance of commercially available anti-RFC1 antibodies, we were unable to demonstrate a significant effect of the repeat expansion on RFC1 protein. Furthermore, the observation of an increased sensitivity of CANVAS cell lines to DNA damage was limited to testing of platinum-based drugs. A more comprehensive study assessing multiple types of DNA damage is therefore warranted to elucidate the intermediate steps linking reduced *RFC1* expression to DNA damage accumulation and subsequent neurodegeneration.

## METHODS

### Determination of the propensity of the AAGGG repeat to form G-quadruplexes

Circular dichroism (CD) spectra were acquired on a J-1500 Spectropolarimeter (Jasco Corporation, Easton, MD, USA) equipped with a Peltier temperature controller set at 20°C. The measurements were performed in a 1 cm quartz cuvette using synthetic oligonucleotides 5’-(AAAAG)_8_-3’ and 5’-(AAGGG)_8_-3’. They were diluted to a final concentration of 1 µM in 10 mM lithium cacodylate buffer at pH 7.2, with 100 mM of LiCl, NaCl, or KCl. Samples were annealed by heating to 95°C for 5 minutes, followed by gradual cooling to room temperature over 4 hours. CD spectra were recorded over the 210 to 340 nm wavelength at a scan speed of 200 nm/min, with a 2s response time and a 1 nm bandwidth. Each spectrum represents the average of four scans and was baseline-corrected to account for contributions from the buffer and chloride salts. CD signals were expressed as Δε (M ¹cm ¹) as a function of wavelength, calculated using the formula Δε = / (32980 × C × l), where represents the observed ellipticity in millidegrees, C is the DNA molar concentration, and l is the path length in centimeters (1 cm). For CD-melting experiments, the ellipticity was monitored at 220 nm (5’-(AAAAG)_8_-3’) or 264 nm (5’-(AAGGG)_8_-3’) in 0.5°C intervals, with a 1°C/min heating ramp from 20°C to 95°C. Before measurement, the samples were equilibrated at 20°C for 5 minutes. Melting temperatures (Tm) were calculated as the average of two independent experiments, using the Van’t Hoff equation, assuming a two-state folded-unfolded G4 transition and equivalent heat capacities of the folded and unfolded states.

Principal Component Analysis (PCA) of CD spectra was performed using spectra from both repeat sequences in the three salt conditions, alongside reference spectra from experimentally resolved nucleic acid structures obtained from (Del Villar-Guerra et al. 2017). PCA was performed using the Circular Analyses Tool (CAT) application in R on mean-centered data.

The first two principal components (PC1 and PC2) explained 40.4% and 33.3% of the variance, respectively, and were plotted to visualise clustering patterns of pathogenic and non-pathogenic repeats relative to G4 and non-G4 reference structures.

### Generation of repeat-containing plasmids by recursive directional ligation

Plasmids containing a range of increasing sizes of CCCTT or TTTTC repeats were generated using recursive directional ligation, as previously described (McDaniel 2010). The method used is graphically illustrated in **Supplementary Fig. 13.** In brief, complementary DNA oligonucleotides containing a sequence of 12 repeated CCCTT or CTTTT units (Sigma-Aldrich), flanked 5’ by Esp3I and BamHI recognition sites and 3’ by PaqCI and NotI recognition sites, were resuspended in Nuclease-free water, mixed in equimolar concentration and annealed by placing the equimolar mix at 95°C for 5 minutes and then allowing for slow cooling at room temperature. Annealed oligos were then ligated into the vector pBluescript II SK+ using by BamHI and NotI (NEB) overhangs through T4 ligase (NEB). Oligomerization of repeat unites was achieved by recursive directional ligation by digesting the plasmids in two separate reactions, one with Esp3I and BsaHI and one with PaqCI and BsaHI (all NEB) to release two fragments of the plasmid, including one containing the entire CCCTT or CTTTT repeat. The double digestion products were separated by gel electrophoresis (1.2% agarose) and the repeat containing fragments were identified based on the expected size compared to a 100bp DNA ladder (New England Biolabs) and purified from the gel (QIAquick Gel Extraction Kit). Ligation of the two repeat-containing fragments was performed with T4 Ligase (NEB) with a 1:1 or 1:2 vector to insert ratio overnight at room temperature and transformed into One Shot Stbl3 Chemically Competent E. coli cells (Thermo-Fisher). Repetition of the same round of recursive directional ligation was used to generate plasmids containing CCCTT or CTTTT repeated sequences of increasing size. Repeat unites were joined seamlessly without interruption. All plasmids were subjected to long-read sequencing to confirm exact repeat size as reported in **Supplementary Table 5**

### *In vitro* transcription assay

Prior to transcription, 1 µg of pBluescript containing 12, 24, 48 and 60/70 CCCTT or TTTTC repeats was linearized with NotI-HF (NEB). Transcription was performed using a HiScribe™ T7 Quick High Yield RNA Synthesis Kit (NEB) and incubated at 37 °C for 1 hour. Aliquots were removed at 30 minutes after the addition of T7 RNA polymerase. 5 µl of transcription product were denatured by adding 5 µl of Invitrogen™ Novex™ TBE-Urea Sample Buffer (2X) and heated at 95°C for 5 min. Samples were separated on a 6% TBE-Urea denaturing gel (Novex, ThermoFisher) at 200V for 1 hour. The gel was post-stained with SYBR™ Gold Nucleic Acid Gel Stain (Invitrogen) and imaged with ChemiDoc imaging system (Bio Rad). Quantification of band intensity was performed with ImageLab Software. Statistical significance was calculated using Brown-Forsythe and Welch ANOVA test with Dunnett’s T3 multiple comparisons post-hoc test.

### Gene reporter vectors

pCMV-Luc Firefly Luciferase reporter vector (TR30004) plasmid was purchased from Origene while pGint plasmid was a kind gift from Dr. Mariano Garcia-Blanco (Addgene plasmid # 24217). The repeat insert was subcloned from pBluescript SK+ into the recipient plasmids using standard restriction enzyme digestion and ligation techniques. Briefly, Firefly Luciferase plasmid and the repeat containing pBluescript vectors were digested with XbaI and NotI restriction enzymes (NEB), while pGint plasmid and pBluescript vectors were digested with NotI and BamHI restriction enzymes (NEB) at 37°C for 1 hour. The restriction digestion products were then run on a TAE agarose gel (1.2%), purified from the gel using QIAquick Gel Extraction Kit (Qiagen).

The two fragments were then ligated at a 1:7 ratio with T4 DNA ligase (NEB) at 16°C overnight, following the manufacturer’s instructions. The subcloned plasmids were then used to transform One Shot Stbl3 Chemically Competent *E. coli* (Invitrogen) following the manufacturer’s instructions, and colonies were grown overnight on Lennox LB agar plates and selected with 50 µg/mL kanamycin for pGint and 100 mg/mL ampicillin for Firefly Luciferase. Colonies were picked and grown overnight in liquid LB Lennox broth. The plasmids were then extracted using QIAprep Spin MiniPrep Kit and HiSpeed Plasmid Midi Kit (Qiagen). All plasmids were subjected to long-read sequencing to confirm correct insertion of the CCCTT or TTTTC repeat containing fragment and to provide exact repeat sizing as reported in **supplementary Table 4.**

### Luciferase and fluorescence assays on reporter vectors

HEK293T cells were cultured in Dulbecco’s Modified Eagle Medium (DMEM) (ThermoFisher Scientific) enriched with 10% fetal bovine serum (FBS) (ThermoFisher Scientific). One day before transfection, HEK293T cells were trypsinised and plated in 96-well white tissue culture plates (Greiner), at a seeding density of about 1×10^4^ cells per well. 50ng Firefly Luciferase plasmids were co-transfected in a 1:1 ratio with a control reporter vector constitutively expressing Renilla Luciferase (Origene), using Lipofectamine^TM^ 3000 Transfection Reagent (ThermoFisher Scientific). After 48 hours, the Dual-Glo® Luciferase Assay System was used to detect firefly and Renilla Luciferase activities in a single well, following the manufacturer’s instructions. Briefly, Dual-Glo® Luciferase Assay Reagent was added to the plate in a 1:1 ratio compared to the volume of the growth medium and incubated in the dark at room temperature for 10 minutes. The Firefly luminescence was measured using FLUOStar Omega. Dual-Glo® Stop & Glo® Reagent was then added to the plate in a 1:1 ratio compared to the initial volume. After 10 minutes incubation in the dark at room temperature, a second measurement of the Firefly Renilla luminescence was obtained. The ratio of Firefly:Renilla luminescence was calculated for each well. All samples were run in triplicates.

For pGint splicing vector experiments, 10 ng pGint plasmids were co-transfected in a 1:10 ratio with a reporter vector constitutively expressing mCherry (mCherry—hFUS) using Lipofectamine^TM^ 3000 Transfection Reagent.

Enhanced green fluorescent protein (EGFP) fluorescence observation was performed using the Spark® multimode multiplate reader at an excitation wavelength of 487 nm and an emission wavelength of 513 nm using recommended exposure time. mCherry red fluorescence observation was performed immediately after at an excitation wavelength of 585 nm and an emission wavelength of 610 nm. The EGFP:mCherry fluorescence ratio was calculated for each well. Non-transfected cells were also observed to evaluate the autofluorescence. All samples were run in triplicates. Statistical significance was calculated using multiple unpaired t-test with Welch correction. Adjusted P-values were calculated using the Bonferroni correction for multiple comparisons.

### Brain donors

Brain donors with a genetically confirmed CANVAS were recruited by the Queen Square Brain Bank (UCL, London, UK), the London Neurodegenerative Diseases Brain Bank (King’s College, London, UK), the Newcastle Brain Tissue Resource (Newcastle, UK), the New York Brain Bank at Columbia University (New York, US), and the Harvard Brain Tissue Resource Center (Boston, US), according to the regulations approved by the local ethics committees. Each centre provided control samples to address for site-specific variability (e.g. sampling and fixation methods, storage conditions).

When available, fresh frozen brain tissue from frontal cortex and cerebellar hemispheres was utilized for expression analysis. Formalin-fixed, paraffin-embedded slides were used for immunohistochemistry.

### Generation of induced pluripotent stem cells from CANVAS patients

Induced pluripotent stem cells (iPSC) lines from two CANVAS patients and healthy subjects were generated by Oxford StemTech (Oxford, UK) from human whole blood derived peripheral blood mononuclear cells (PBMCs) or human skin fibroblasts **Supplementary Table 3.** Local ethical committee approval and informed consent were obtained prior to human specimen sampling. Skin fibroblasts were reprogrammed into iPSC via Cytotune Sendai virus iPSC reprogramming system. Blood samples were processed for PBMC isolation and cryopreservation. All PBMC lines were exposed to erythroblast derivation media and growth factors. Eritroblasts were subjected to Sendai virus-based iPSC reprogramming using the company’s ReproPlex™ technology. iPSC colonies were manually isolated and expanded to derive 2 iPSC lines per donor and frozen between passage number 5-7. Flow cytometry of markers NANOG and TRA-1-60 was performed to confirm pluripotency in reprogrammed lines. Molecular karyotyping with Illumina’s Infinium Global Screening Array was performed in donor PBMCs and reprogrammed lines to confirm genomic integrity. All lines were confirmed to be Mycoplasma free using Lonza’s MycoAlert kit. iPSCs were grown on Geltrex-coated wells and maintained in mTeSR Plus (StemCell).

### Generation of Isogenic iPSC Lines

Isogenic iPSC lines were generated by CRISPR/Cas9-mediated excision of most or all of the AluSx3 element containing the pathogenic repeat expansion in intron 2 of *RFC1*. Using Lipofectamine Stem, cells were co-transfected with TrueCut HiFi Cas9 Protein (Invitrogen) and two chemically modified sgRNAs (Synthego) (Sequences in **Supplementary Table 6).** For both patient lines the same 3’ sgRNA was used (L2), whereas the 5’ sgRNA differed between lines: sgRNA M1, cutting 36nt inside the AluSx3 element, was used for the CANVAS1 line and sgRNA R3, cutting 7nt outside the AluSx3 element, was used for the CANVAS2 line. After single-cell sorting, isolated clones were genotyped using a dual PCR strategy using flanking and internal primers as shown in **Supplementary Fig. 3A-B** and **Supplementary Table 7.** Additionally, edited lines were assessed using repeat-primed PCR using established protocols (Cortese et al. 2019) (**Supplementary Fig. 3C)**. Successful editing **(Supplementary Fig. 3D)** and Genomic integrity (**Supplementary Fig. 3D)** of edited lines was confirmed using low-pass Nanopore long read whole genome sequencing at UCL Institute of Neurology long-read facility.

### Differentiation of iPSC into sensory neurons

Using Lipofectamine Stem (Invitrogen), iPSCs were transfected with vectors EF1a-Transposase and PB-tetO-NB (Addgene #197092) and then selected with Puromycin 0.5ug/mL for at least 14 days before being cryopreserved.

Piggybac-integrated iPSC lines were differentiated to sensory neurons via doxycycline-inducible forced expression of Neurogenin-2 and Brn3a (POU4F1) transcription factors. Briefly, iPSCs were dissociated with Accutase and 1.0×10^5^ cells were resuspended in Neuronal Differentiation Medium (NDM) supplemented with 50nM Chroman1 (MedChemExpress) and 2mg/mL doxycycline (Sigma Aldrich), and seeded onto a Geltrex-coated T175 flask. After 48 hours, induced neurons were dissociated using Accutase, counted and cryopreserved. Upon thawing, neurons were seeded onto plates double coated with 0.1% Polyethylenemine (Sigma) and Geltrex (Gibco) in NDM supplemented with 20uM Bromodeoxyuridine (BrdU) and 2mg/mL doxycycline. On days 4,6 and 8, full media changes were carried out without BrdU. Neutrophic factors NT3, BDNF, bNGF and GDNF (all at 10ng/mL) were added from day 8, and half media changes were performed every 2 days.

### Differentiation of iPSC into lower-motor neurons

Unedited and isogenic iPSC clones were differentiated into motor neurons following a 14-day protocol, based on the addition of small molecules and as described in (Pagliari et al. 2025). On day 0, iPSCs were dissociated with Accutase (Stemcell Technologies) and resuspended in a N2B27-based differentiation medium (1:1 DMEM/F12 - Neurobasal medium, N2- supplement, B27, 2 mM L-glutamine, 1% Penicillin-Streptomycin, 0.1 mM β-mercaptoethanol; all from ThermoFisher). Dual SMAD inhibitors LDN-193189 (0.1 μM) and SB-431542 (20 μM) were supplemented from day 0 together with the Wnt inhibitor CHIR-99021 (3 μM), potently inducing the neuralization of cells and the formation of embryoid bodies (EBs). The addition of retinoic acid (100 nM) on day 2, of smoothened agonist (SAG, 500 nM) on day 4, and of DAPT (10 μM) on day 9 promoted the differentiation towards spinal motor neurons. On day 7 the SMAD inhibitors and CHIR-99021 were removed. EBs were dissociated between day 11 and 14 and replated on Geltrex-coated 6-well plates or 96-well Phenoplates (PerkinElmer). Post-mitotic neurons were maintained in a N2B27-based maintenance medium supplemented with 100 nM RA, 500 nM SAG, 10 μM DAPT (all withdrawn on day 7 after replating) and the neurotrophic factors BDNF, GDNF, and CNTF (10 ng/mL; Peprotech). Half-medium change was performed every 3-4 days and experiments were performed between day 18 and day 25 of differentiation.

### Assessment of maturation markers in iPSC-neurons

At day 14 (sensory neurons) or 20 (motor neurons), cells were washed with PBS and fixed with 4% paraformaldehyde for 15 min. Fixed cells were permeabilized with PBS-Triton-X 0.25% for 20 min, subsequently washed with PBS and incubated with 5% goat serum in PBS-Tween 0.1% (Thermo Fisher) for 1 h at room temperature. Primary antibodies were applied overnight at 4°C. iPSC-motor neurons were stained for the markers HB-9, TUBB3, MAP-2; iPSC-sensory neurons were stained for BRN3A and TrkB.

Cells were then washed three times and incubated with fluorescent conjugated secondary antibody (Alexa Fluor® 488 and Alexa Fluor® 647) for 1 h, followed by nuclear counterstaining with DAPI for 15 min. Cells were imaged using the Opera Phenix^®^ High-Content Screening System (PerkinElmer Inc.), equipped with the Harmony software.

Morphological properties (size and roundness) were defined for post-mitotic neurons, as defined by a positive signal for maturation markers, and used to identify and filter out contaminating cells.

### Assessment of *RFC1* mRNA expression by quantitative RT-PCR

For iPSC-derived motor and sensory neurons, cells were harvested in 1X RNA/DNA Shield (Zymo). RNA was extracted using the Zymo Quick-RNA Miniprep kit, according to manufacturer’s instructions. For brain samples, total RNA for each sample was extracted by Bioexpedia (Aarhus, Denmark) from 80 mg of bulk tissue of frontal cortex and cerebellar hemispheres.

cDNA was synthesised from 250 ng (brain) or 500 ng (iPSC-neurons) of RNA using the Maxima H Minus cDNA Synthesis Mastermix (ThermoFisher).

For Real-Time quantitative PCR, 4 uL of cDNA samples were mixed with 4 uL of TaqMan Fast Advanced Master Mix, 1 uL of TaqMan probes (RFC1 Hs01099126_m1; TATA-binding protein (TBP) (Hs00427620_m1)), and 1 uL of nuclease-free water. Samples were then run on QuantStudio 7 Flex Real-Time PCR System (Applied Biosystems) for 20 sec at 95°C, followed by 40 cycles of 1 sec at 95°C and 20 sec at 60°C. TBP was selected as the housekeeping gene, as its transcript abundance in neurons lies within the range anticipated for RFC-complex. Relative gene expression was analysed using qRAT (Flatschacher, D., Speckbacher, V. & Zeilinger, S. qRAT: an R-based stand-alone application for relative expression analysis of RT-qPCR data (Flatschacher et al. 2022).

Data were presented as normalized fold expression change (ΔΔCT) and plotted using GraphPad Prism 7 software (GraphPad Software).

### RNA-seq in brain tissue and iPSC-neurons

Bulk RNA-seq was performed on brain tissue (cerebellar hemispheres and frontal cortex from CANVAS patients, N=5 and N=6, respectively, and healthy controls, N=4 and N=5, respectively), and iPSC-sensory neurons (two isogenic lines and including CANVAS unedited (AluSx3_exp_ +/+); heterozygous corrected (AluSx3_exp_ +/-), and homozygous corrected (AluSx3_exp_ -/-) , one line with 3 replicates (CANVAS1) and one with 2 replicates (CANVAS2);

For all samples, libraries were prepared at UCL Genomics from 250ng of total RNA using the Watchmaker RNA Library Prep kit with Polaris Depletion (Watchmaker Genomics) according to manufacturer instructions. High yield, adaptor-dimer free libraries were confirmed on Agilent TapeStation 4200 using the High Sensitivity DNA 1000 assay (Agilent) and normalised to 10nM. Libraries were pooled and sequenced on the NextSeq 2000 instrument (Illumina) at 800pM, using a 57bp paired-end run with corresponding 8bp unique dual sample indexes and 8bp unique molecular index.

For brain samples, sequencing data were analysed with the nf-core/rnaseq pipeline v3.14.0, which includes FastQC v0.12.1 for quality control and STAR v2.7.9 for spliced alignment.

Subsequently, reads were processed with featureCounts v2.0.1 and Gencode v35 to obtain gene expression count. Principal Component Analysis (PCA) and differential gene expression (DGE) analysis were performed with DESeq2 including RIN and sex as covariates in the model. For the functional characterization of differently expressed genes, we used the Gene Ontology Enrichment Analysis method included in R package clusterProfiler (function “enrichGO” applied to Biological Process ontology, with FDR <0.05). To reduce the number of redundant terms, the “simplify” method (also included in clusterProfiler) was applied with cutoff=0.5. Differential exon usage analysis was performed with DEXSeq, including exons from all available transcripts annotated in Ensembl v111.

For iPSC-sensory neurons, sequencing data were analysed with nf-core/rnaseq as described above. Read counts were obtained with salmon v1.10.3.

PCA, DGE analysis and Gene Ontology analysis were performed as described above. For DGE, batch and parent line were included as covariates.

### Amplicon-based targeted long-read sequencing of RFC1 full transcript

Full length RNA-seq was performed on iPSC motor neurons (CANVAS n=3 and controls n=3) and sensory neurons (isogenic cell lines described above, one replicate each). cDNA was synthesized using random hexamers and SuperScript III Reverse Transcriptase (Invitrogen). Full-length transcripts were PCR amplified from cDNA in 50uL PCR reactions of Phusion High-Fidelity PCR Master Mix (Thermo Scientific) with primers designed to anneal on the 5’ (5’-GGATCCTGAGCCTCGATAACAG – 3’) and 3’ UTR (5’-TTAGTACTACTGCTCCCAGGCT – 3’) of the *RFC1* MANE transcript (<u>NM_002913.5</u>).

Amplicons were bead purified and nanopore sequenced as PCR libraries by Full Circle Labs (London, UK). Sequencing reads were mapped to GRCh38 using minimap2. Transcript usage was quantified and plotted with Bambu v3.8.3, using standard options (including filtering of transcripts with total read count > 2). Differential Transcript Usage analysis was performed with DEXSeq as described in Bambu reference manual.

### Knockdown of RFC1 in sensory neurons and continuous live-cell imaging

Cryopreserved day 2 induced sensory neurons were seeded onto 96-well PhenoPlates (Revity) at a density of 4.0 x 10e4 cells/well. On day 14, wells were transduced in triplicate with lentivirus expressing either RFC1 shRNA-mCherry (cat. VB240716-1228rae [VectorBuilder], target sequence ATACTCACTTCAAGCTATAAA) or scramble shRNA-mCherry (cat. VB010000-0015hpw [VectorBuilder], target sequence CCTAAGGTTAAGTCGCCCTCG) at a predicted MOI 1 and continuous live-cell imaging was started on a Incucyte S3 (Sartorius) microscope inside a 37°C 5% CO2 incubator. For each well, 9 images at 20X magnification were obtained every 8 hours, for both phase-contrast and red fluorescence channels. On day 15, lentivirus-containing media was replaced with fresh media. Thereafter, half-media changes were performed every 3-4 days. Virtually every cell in both conditions was confirmed to develop intense mCherry+ fluorescence 96hrs post-transduction. Segmentation of live neurons was performed in phase-contrast images using the following parameters in Incucyte software version 2023A: Object diameter 10.0uM, Threshold Sensitivity 4, Texture Sensitivity 4, Edge Sensitivity 8, Adjust size 2px, Minimum area 180, Maximum area 400. Objects meeting these criteria were termed ’live neurons’, and their counts for each image were exported and analysed on GraphPad Prism version 10. Baseline (100%) was defined as the mean of the first four recordings for each individual image. Mean and SEM were calculated and plotted from 27 individual images for each condition. Statistical significance was calculated using a mixed-effects model with Geisser-Greenhouse correction and P-values were adjusted using the Bonferroni method for multiple comparisons.

### Quantification of apoptotic markers after cisplatin or oxaliplatin continuous treatment in iPSC neurons

Cisplatin (Sigma) and oxaliplatin (Sigma) stock solutions were prepared as described above in 0.9% NaCl at a concentration of 4 mM, sterilized by filtration and kept at 20°C. Working dilutions were prepared in complete RPMI medium. Untreated cells were exposed to the same NaCl concentration of drug exposed samples to harmonize culture conditions.

On day 7 (sensory neurons) or 20 (motor neurons) of differentiation, cells were treated with different concentrations of cisplatin (0,5, 10 uM) for 24 hours. Three technical replicates were used for each condition. At the end of the treatment, medium was completely removed and cells were fixed in PFA 4%. Immunofluorescence was performed as described for the assessment of neuronal maturation markers. Neurons were stained for cleaved Caspase-3 and Tuj1. After the staining, cells were imaged using the Opera Phenix^®^ High-Content Screening System (PerkinElmer Inc.), equipped with the Harmony software. Images were captured using a 40x water immersion lens and settings were optimized for each channel. For each well, nine pre-defined fields of view and 18-22 Z-stacks were acquired. Analysis involved maximum projection of Z-stacks, using nuclei detection algorithms with custom settings for noise reduction and nuclear segmentation. To quantify the number of apoptotic cells, a threshold for AlexaFluor 488 intensity was manually defined to exclude background noise. Apoptotic rate was measured as percentage of positive cleaved Caspase-3 nuclei out of the total neuronal cells. Statistical significance was calculated using Brown-Forsythe and Welch ANOVA test with Dunnett’s T3 multiple comparisons post-hoc test.

### Sensitivity to platinum-based DNA damaging agents in LCLs

B Lymphoblastoid cell lines (B-LCLs) of CANVAS patients and age-matched healthy controls were generated at the European Collection of Authenticated Cell Cultures (ECACC) by Epstein-Barr Virus (EBV) transformation of peripheral blood lymphocytes from CANVAS patients and healthy controls. Local ethical committee approval and informed consent were obtained prior to human specimen sampling. Cells were grown in suspension in RPMI medium supplemented with 15% fetal bovine serum, 2mM L-Glutamine, 100 U/mL penicillin and 0.1 mg/mL streptomycin at 37°C in a 5% CO2 atmosphere.

Cisplatin (Sigma) and oxaliplatin (Sigma) stock solutions were prepared as described above in 0.9% NaCl at a concentration of 4 mM, sterilized by filtration and kept at 20°C. Working dilutions were prepared in complete RPMI medium. Untreated cells were exposed to the same NaCl concentration of drug exposed samples to harmonize culture conditions.

For the evaluation of cell sensitivity, 5×10^5^ cells/mL were treated with different concentrations of Cisplatin (ranging from 3 μM to 30 μM) or Oxaliplatin (from 1 μM to 5 μM) for 24 h and 48 h. At each time point, cell viability was measured through RealTime-GloTM MT Cell Viability Assay (Promega) based on a non-lytic and metabolic luciferase reaction, following manufacturer’s instructions. Briefly, an aliquot of cells (40 μL) was mixed with equal volume of 2X RealTime-GloTM Reagent and spotted in a 384-well plate. After incubation for 1 h at 37°C, 5% CO2, luminescence was measured through the GloMax® Discover microplate reader (Promega). Statistical analysis was performed with GraphPad Prism 9.0. Unpaired parametric t-test with Welch correction was performed to compare two groups assuming unequal variance between the two. P-value<0.05 was considered statistically significant.

### Quantification of apoptotic markers after cisplatin or oxaliplatin continuous treatment in LCLs

Patients and controls-derived LCLs were treated with different concentrations of platinum-derivative agents as described above. After treatment, cells were pelleted and resuspended in RIPA lysis buffer (1% NP40, 50 mM TrisHCl pH 8.0, 150 mM NaCl, 0.1% SDS, 0.1% DOC, 1X protease inhibitor cocktail (Sigma), 1X phosphatase inhibitor cocktail (Roche Diagnostics)) for 30 min in ice, sonicated and centrifuged (13’000 g, 5 min). Total protein content of each sample was quantified using a BCA assay (Thermo Fisher Scientific). Whole cell lysates (20-25 μg) were incubated with loading buffer (300 mM Tris-HCl pH 6.8, 10% SDS, 40% glycerol, 600 mM DTT, 5% 2-mercaptoethanol, and 0.2% bromophenol-blue) and denatured for 5 min at 95°C. Samples were separated on a 4-20% gradient SDS-polyacrylamide gel (Bio-Rad) in 1X Tris/glycine/SDS running buffer and transferred onto a nitrocellulose membrane using Turbo Transfer Pack (Bio-Rad) in the Trans-Blot Turbo Transfer System (Bio-Rad). Membranes were blocked for 1 hour in 6% skimmed milk and incubated overnight at 4°C with 2.5% skimmed milk with the following antibodies recognizing human proteins: anti β-actin (2Q1055) mouse monoclonal (cat. no. sc-58673, 1:1’000, Santa Cruz biotechnology), anti p53 (DO1) mouse monoclonal (cat. no. sc-126, 1:1’000, Santa Cruz Biotechnology), anti-cleaved Caspase 3 (D175), rabbit polyclonal (cat. no. 9661S, 1:1’000, Cell Signaling) and anti γH2A.X (JBW3) mouse monoclonal (cat. no. 05-636, 1:1’000, Sigma). Primary antibodies were probed by a secondary horseradish peroxidase-conjugated antibody (antimouse, cat. no. 115035146; antirabbit, cat. no. 111035144, 1:3’000, Jackson Immuno Research Laboratories) for 1 h at RT. Protein bands were visualized using an enhanced chemiluminescent reagent (Westar Eta C Ultra 2.0, Cyanagen) through ChemiDoc XRS (Bio-Rad). Bands were quantified using QuantityOne software (BioRad), corrected for the intensity of the corresponding band obtained with the anti β-actin and normalized to the appropriate control sample. Western blotting experiments were repeated at least three times and with independent biological replicates. Statistical significance was calculated using Brown-Forsythe and Welch ANOVA test with Dunnett’s T3 multiple comparisons post-hoc test.

### Assessment of cell cycle and apoptosis after cisplatin or oxaliplatin pulse treatment in LBLs

To monitor cell cycle progression, 2.5×10^5^ cells/mL were treated with 10 μM or 30 μM cisplatin for 1h, then washed in PBS and let recover in complete medium for 24 h and 48 h at 37°C, 5% CO2. At each time point (0 h, 24 h and 48 h of recovery) cells were pulsed with 30 µM BrdU (Sigma) for 30 min, washed twice with PBS, fixed in ice-cold 70% ethanol and stored at 4°C overnight. DNA was denatured with 2N HCl/0.5% Triton X-100 and neutralized with 0.1M sodium borate pH 8.5. After centrifugation (2’500 g, 10 min, 10°C) the cell pellets were resuspended for 1h at room temperature (RT) with a mouse monoclonal anti-BrdU antibody (cat. no. BD347580, 1:100 in 1% BSA with PBS 0.5% Tween-20, Becton Dickinson). Cells were then centrifuged (2’500 g, 10 min, 10°C) and the pellets were resuspended for 1h at RT with fluorescent antibody Alexa anti-mouse-488 (cat. n. ab150113, 1:500 in 1% BSA with PBS 0.5% Tween-20, Abcam). Samples were then incubated with 10 µg/mL Propidium Iodide and analysed on a S3 Flow Cytometer (Bio-Rad), at least 10’000 events were recorded, and data were examined using FCS Express software (DeNovo).

The FITC Annexin V/Dead Cell Apoptosis Kit (Invitrogen) was used to detect early and late apoptosis in cells following manufacturer’s instruction. Briefly, 2.5×105 cells/mL were treated with 5 μM or 10 μM cisplatin for 24 h, then washed with PBS and resuspended in 1X binding buffer (10 mM HEPES–NaOH pH 7.4, 144 mM NaCl, 25 mM CaCl2) at the concentration of 106 cells/mL. For each sample, 100 μL of cell suspension were incubated with Annexin V-FITC (0.2 µg/µl) and Propidium Iodide (0.05 µg/µl), incubated in the dark for 15 min at RT and immediately analysed on a S3 Flow Cytometer (Bio-Rad). At least 10’000 events were recorded, data were examined using Pro-Sort software (Bio-Rad).

### Dot blotting for quantification of platinum-(GpG) DNA adducts in LCLs

LCLs were seeded at 5 × 10 cells/mL in 6-well plates and treated with 30 µM cisplatin for 1.5 hours. After treatment, cells were collected with no recovery (t=0) or recovered in complete medium for 6, 24, 48 and 72 hours (t=6, t=24, t=48, t=72) at 37°C, 5% CO2. At each time point, cells were centrifuged (2’000 rpm, 5 min, RT), washed with 1mL of 1x PBS (EuroClone SpA, Milan, Italy) and centrifuged again (5’000 rpm, 10 min, 4°C). Cell pellets were then used to obtain genomic DNA (gDNA); extractions were performed with NucleoSpin Tissue kit (Macherey-Nagel, Dueren, Germany) according to manufacturer’s instruction. DNA integrity and quantification were measured through a spectrophotometric method. For each time point, 2µg of gDNA was diluted in water up to 50µL, denatured by heating (10 min, 95°C) and immediately cooled on ice. Subsequently, 50µL ice-cold ammonium acetate (Ci=2M, Cf=1M) was added and samples vortexed for 10 sec. Using a dot-blot apparatus (Bio-Rad, Hercules, CA, USA), 100µL of each sample were transferred on Hybond-N+ membrane (Cytiva, Amersham, UK) pre-soaked in 1M ammonium acetate for 2 min. Thereafter, the membrane was washed with ammonium acetate 1M for 2 min, neutralized with 6x sodium saline citrate for 5 min and rinsed in water for 1 min. DNA was fixed onto the membrane by cross-linking (Stratagene, La Jolla, CA, USA). The membrane was blocked in 5% skimmed milk/TBS 1x for 1 h at RT and incubated with an antibody specific for Pt-d(GpG) DNA crosslinks (courtesy of Prof. J. Thomale/ONCOLYZE, Essen, Germany) at a dilution of 1:250 (in skimmed milk 3%/TBS 1x) for 1 h at RT. The primary antibody was detected with an anti-rat HRP-linked secondary antibody (1:10’000) for 1h at RT. The immunoblotting was revealed using enhanced chemiluminescent reagent (Westar Eta C Ultra 2.0, Cyanagen, Bologna, IT) through ChemiDoc XRS (Bio-Rad, Hercules, CA, USA). Dots were quantified using ImageJ software (NIH, Bethesda, MD, USA). Signal intensities were corrected with signals obtained with anti-single-stranded DNA antibody and normalized toward control sample (untreated). Statistical analysis was performed with GraphPad Prism 9.0 software on at least three biological replicates (unless otherwise stated). Unpaired t test with Welch correction was performed to compare two groups assuming unequal variance between the two.

### H2A.X immunostaining and quantification on post-mortem brain tissue

Formalin-fixed paraffin-embedded (FFPE) slides of brain tissues were immersed in xylene and then hydrated in graded ethanol solutions. All the staining were carried out on the automated Ventana Discovery Ultra machines (Roche). Heat-induced epitope retrieval was performed, using Tris-EDTA buffer (pH=7.8) heated to 95 °C. Sections were incubated with the primary antibody (rabbit anti-γH2A.X, ab81299, Abcam) at a dilution of 1:250 for 4 hours. Following incubation for 1 hour with a goat anti-rabbit secondary antibody (ab207995, Abcam), antigen detection was performed using 3,3’-diaminobenzidine (DAB, Roche 760-124). Cells were finally counterstained with haematoxylin (Roche, 760-2021) and bluing reagent (Roche, 760-2037).

Slides were scanned at 40× magnification on a Hamamatsu NanoZoomer S360 by the UCL IQPath Science and Technology Platform. The whole slide images (WSI) were made available on NZConnect (Hamamatsu), a web-based image viewing platform. Annotations for the Purkinje Cell layer (PL) and granular layer (GL) were manually drawn on NZConnect.

These annotations and WSI were imported into QuPath v0.5.1 for image analysis(Bankhead et al. 2017). Cell segmentation was performed within the PL band and GL annotations using Cellpose-SAM (Pachitariu et al. 2025) via the QuPath Cellpose extension (available from: https://github.com/BIOP/qupath-extension-cellpose). A background-corrected DAB mean intensity value was calculated by subtracting the background intensity from the mean DAB intensity of each detection object. Non-cellular detection objects were classified as those with mean haematoxylin intensity <0.1 and mean background-corrected DAB intensity <0.25. γH2A.X positive cells were classified as the remaining detection objects with mean background-corrected DAB intensity ≥0.1. The percentage of γH2A.X positive cells was calculated with the following formula: γH2A.X positive cells/Total cells × 100. Statistical significance was calculated using unpaired t-test with Welch correction.

### *Drosophila* stocks and experimental procedures

Unless otherwise specified, flies were kept on standard fly food at 25°C in 12 h light: 12 h dark cycles. The following fly stocks were obtained from the Bloomington *Drosophila* stock center: *actin*-GAL4 (y[1] w[*]; P{Act5C-GAL4-w}E1/CyO, BDSC #25374) and nSyb-GAL4 (y[1]w[*]; P{w[+m*]=nsyb-GAL4.S}3, BDSC #51635) were used to drive RNA interference (RNAi) expression ubiquitously and in post-mitotic neurons respectively. Both driver insertions were outcrossed into an isogenized background (iso31) for five generations, with the X-linked y[1] marker removed in the process. The *Gnf1* UAS-shRNA expressing line BDSC #35423 (y[1] sc[*] v[1] sev[21]; P{y[+t7.7] v[+t1.8]=TRiP.GL00346}attP2/TM3, Sb[1]) was used to knock down *Gnf1* expression *in vivo*.

### Quantification of *Drosophila* Gnf1 expression after neuronal-specific knockdown

Total RNA was extracted from dissected brain tissue of *nsyb* > *Gnf1* RNAi and *nsyb* > *mCherry* RNAi (control) 3-7 day old adult flies using phenol chloroform extraction method. RNA concentration and purity of the samples were assessed using NanoDrop equipment (NanoDrop Technologies Inc., Wilmington, DE). 0.5 µg total RNA was reverse transcribed (Superscript III, Thermofisher). Technical triplicates of the qPCR samples were prepared as a 10 µL approach with the Fast SYBR™ Green Master Mix (Thermofisher), 500 nM of each primer (forward: CAACGCGGCATTGACTCCT; reverse: CGTCTCTCCACTTTTGGCCTC), and 1 µl of the reverse transcription reaction. qPCR was performed using the QuantStudio 3 Real-Time PCR System (Thermo Fisher Scientific) equipped with QuantStudio Design&Analysis Software v1.4.3 (Thermo Fisher Scientific).

The PCR conditions included a pre-run at 95°C for 5 min, followed by 40 cycles of 30 s at 95°C, 30 s at 58°C and 45 s at 72°C. PCR amplification specificity was determined by melting curve analysis with a range from 60°C to 95°C. The values of the cycle threshold (CT) of the target mRNAs were normalized to the mRNA of RpL4 (forward TCCACCTTGAAGAAGGGCTA; reverse TTGCGGATCTCCTCAGACTT). For relative gene expression, the comparative cycle threshold (ΔΔCT) values were calculated with the QuantStudio Design&Analysis Software (Thermo Fisher Scientific) with *RpL4* as housekeeping gene and expressed as x-fold change to controls.

### Assessment of DNA damaging agents in *Drosophila*

Cisplatin (Sigma) was dissolved in distilled water at 37°C to achieve stock concentration of 5mM and mixed into 10 ml of fly food at a final concentration of 100 µg/ml. An equal amount of distilled water was mixed into fly food as control.

### *Drosophila* lifespan assays

Lifespan assays were performed on 100 newly eclosed male flies from each experimental and control group, both fed with cisplatin and on standard fly food only. Flies were placed in vials of 10 flies per group and transferred into new vials every 2 days. The number of dead flies were recorded at every transfer and the data were plotted as a Kaplan-Meier survival plot. Statistical significance was calculated using the Mantel-Cox Log rank test.

### *Drosophila* locomotor activity assays

The *Drosophila* Activity Monitor (DAM) was used as previously described to quantify locomotor levels in adult flies (Kratschmer et al. 2021). Fly movement was measured at 25°C across a 24 h period comprised of a 12 h light: 12 h dark cycle. Male flies were collected and loaded into glass behavioural tubes (Trikinetics inc., MA, USA) at the following age groups: 3-7 day old, 21-23 day old and 40-42 day old. Cisplatin treated male flies were recorded by DAM at 10-13 days old only. The behavioural tubes contained 4% sucrose and 2% agar for untreated flies and 4% sucrose and 2% agar with 100ul/ml cisplatin for treated flies. All flies subjected to the DAM were acclimatised to the monitor conditions for 48 h prior to the experimental period. Statistical significance was calculated using the Brown-Forsythe and Welch ANOVA or the Kruskal-Wallis test, depending on the distribution of the data, and followed by Dunnett’s T3 multiple comparisons post-hoc test.

### Quantification of DNA damage in *Drosophila* brains

Brains were dissected in phosphate-buffered saline (PBS) (Sigma Aldrich) at the time point indicated and immuno-stained as described previously(Wu and Luo 2006) Briefly, brains were fixed by 20 min room temperature incubation in 4% paraformaldehyde (MP biomedicals) and blocked for one hour in 5% normal goat serum in PBS+0.1% Triton-X (Sigma-Aldrich) (PBT). Brains were incubated overnight at 4°C in a primary antibody against the phosphorylated form of H2Av (mouse UNC93-5.2.1, Developmental Studies Hybridoma Bank) at a final concentration of 1:200. Brains were then washed three times in PBT and incubated overnight at 4% in secondary antibody: goat anti-mouse AlexaFluor 647 (Life Technologies) at 1:1000. The counterstain was DAPI (ThermoFisher Scientific) at 1:1000. Brains were washed three times, then mounted and imaged in SlowFade Gold anti-fade mountant (ThermoFisher Scientific). Images were taken with a Zeiss LSM 710 confocal microscope with an EC ‘Plan-Neoflar’ 20x air objective, taking z-stacks through the entire brain with step size of 1 μm. Images were analysed using ImageJ: z-stacks were 3-D projected using a maximum intensity projection and regions of interest were drawn around the central brain with exclusion of optic lobes. Mean background fluorescence values were subtracted, and values normalised against DAPI counterstain. Images were compared to controls take on the same day with the same settings and normalised to the means values of controls. Statistical significance was calculated using the Brown-Forsythe and Welch ANOVA test with Dunnett’s T3 multiple comparisons post-hoc test.

### Heterozygous RFC1 expansion and risk of oxaliplatin-induced peripheral neuropathy in patients with cancer

Individuals who underwent oxaliplatin treatment were retrospectively identified through the review of local databases at several hospitals and academic institutions (Toxic Neuropathy Consortium). All participants provided informed consent and the study was approved by local institutional review boards. Demographic and clinical data were collected, including sex, age, number of oxaliplatin cycles and cumulative dosage. Eligible patients were those without a history of neuropathy, no prior exposure to chemotherapy, and who had received a cumulative dose of at least 1000 mg of oxaliplatin. The severity of chemotherapy-induced neuropathy was scored according to the Total Neuropathy Scale – clinical version (TNSc) (10.1111/j.1085-9489.2006.00078.x). A clinically significant neuropathy was defined by a TNSc score >7. All participants received *RFC1* genetic testing by flanking PCR and repeat-primed PCR for the AAGGG expansion. In case of no amplifiable PCR products at flanking PCR, repeat-primed PCR for the non-pathogenic configurations AAAGG and AAAAG was also performed. Heterozygous AAGGG expansions carriers were defined as those presenting an amplifiable, non-expanded product at flanking PCR and a positive repeat-primed PCR for the AAGGG expansion.

Chi-square test was performed to compare the distribution of *RFC1*-carriers between participants with clinically relevant neuropathy vs no/mild neuropathy. Logistic regression was performed to describe the association between the RFC1 expansion and the risk of developing a clinically relevant neuropathy, corrected for age, sex, and centre. P-value<0.05 was considered statistically significant.

## SUPPLEMENTARY FIGURES

Supplementary Figure 1. Pathogenic RFC1 repeats form stable parallel G-Quadruplex structures

**A.** Bar graphs show the estimation of the fractions of the tertiary structural elements of 5’-(AAGGG)8-3’ G-quadruplex in the presence of 100 mM of KCl, NaCl, or LiCl. These data were calculated by exploiting the algorithm developed by *(Del Villar-Guerra et al. 2017)*

**B.** Thermal denaturation profiles of *1 µM* 5’-(AAGGG) -3’ (top row) and 5’-(AAAAG) -3’ (bottom row) in 10 mM lithium cacodylate buffer at at pH 7.2, in the presence of 100 mM KCl (left), 100 mM NaCl (middle), or 100 mM LiCl (right), at 20 °C, monitored by CD. Data points represent experimental measurements, and solid lines correspond to the best-fit curves used to estimate melting temperatures (Tm).

**C.** Principal component analysis of CD spectra from pathogenic 5’-(AAGGG) -3’ and non-pathogenic 5’-(AAAAG) -3’ repeats in the presence of 100 mM KCl, NaCl, or LiCl, alongside reference spectra from experimentally resolved G4-forming nucleic acid structures: parallel (yellow), hybrid (blue), and antiparallel (green) G4s (Del Villar-Guerra et al. 2017). AAGGG repeats (red diamonds) cluster with G-quadruplex references (parallel in K , hybrid in Na /Li ), whereas AAAAG repeats (black diamonds) are clearly separated from the G4 structural clusters, indicating that they do not adopt a canonical G4 topology under the tested conditions. PC1 and PC2 explain 18.9 % and 62.8 % of the variance, respectively.

**Supplementary Figure 2. Transcriptional changes in the cerebellum and frontal cortex of CANVAS patients**

**A. Principal** component analysis (PCA) of gene-level expression data from RNA-seq of CANVAS (n=5) and healthy control (HC, n=4) cerebellar hemisphere samples (**left**) and CANVAS (n=6) and healthy control (HC, n=4) frontal cortex samples (**right**). PCA plotting of the first two PC shows a strong dependence of PC1 on the RNA integrity number (RIN).

**B. Volcano** plot showing differentially expressed (4 upregulated and 1 downregulated) genes in CANVAS vs **control** frontal cortex.

**C.** Normalised read **counts** of for all subunits of Replication Factor Complex (*RFC1-5)* in CANVAS vs control cerebellar hemispheres and frontal cortex. Data are shown as single data points, mean and standard error. The statistical significance was calculated using unpaired t test analysis.

**D. EXSeq** plots of *RFC1* splicing showing similar splicing and alternative exon usage in CANVAS vs controls cerebellar hemispheres and frontal cortex.

**Supplementary Figure 3. RFC1 repeat expansion associated haplotype acts as a cerebellum specific eQTL**

***A.*** *RFC1* region and expansion-associated haplotype. Significant expression Quantitative Trait Loci (eQTLs) according to GTEx dataset are shown as bar charts for different tissues, with Normalised Effect Size (bar colour) and p-value (bar height). The haplotype corresponds to a linkage disequilibrium (LD) block, showed as red dashed triangle on the LD heatmap. (Fig. modified from GTEx website).

**B. RFC1** repeat size and motif in 89 PacBio long-read whole genomes part of the UK 100,000 Genome Project (100KGP). *RFC1* repeat size is stratified according to the haplotype-tagging SNP rs6815219. Each point represents one allele. The haplotype-tagging SNP rs6815219 is associated with expanded repeats, with largest sizes (over 140 repeats) observed for the pathogenic AAGGG motif.

**C.** The **haplotype**-tagging SNP rs6815219 is associated with reduced expression of *RFC1* in Cerebellum (**left and Supplementary Fig. 3A**). Linear regression is overlayed as dashed line. The effect size is larger in carriers of pathogenic AAGGG expansion *vs* non-pathogenic AAAAG expansions (**right**) (reduction of *RFC1* normalised expression in AAGGG_exp_ vs non-expanded =1.11 (Cohen d), p=0.0062, median reduction of *RFC1* normalised expression in AAAAG_exp_ vs non-expanded =0.38 (Cohen d), p=0.025)

**Supplementary Figure 4. Generation and Quality Control of isogenic iPSC lines**

**A. Schematic** representation of RFC1 intron 2 showing the approximate location of 3’ and 5’ sgRNAs used for excising AluSx3 element and its flaking AAGGG expanded repeat (AluSx3_exp_) by CRISPR/Cas9 editing as well as location of primers used for genotyping of edited lines.

**B.** Gel **electrophoresis** of PCR product (**left**) and repeat-prime PCR plots targeting the pathogenic AAGGG repeat (**right**) shows unedited (AluSx3_exp_ +/+), monoallelic corrected (AluSx3_exp_ +/-) and biallelic corrected (AluSx3_exp_ -/-) isogenic iPSC lines from two CANVAS patients.

**C. Visualization** on IGV of reads aligned to the short tandem repeat and flanking AluSx3 element showing the presence biallelic AAGGG expansion in CANVAS1 and CANVAS2 unedited line (AluSx3_exp_ -/-), and of a deletion corresponding to short tandem repeat and flanking AluSx3 from either one (AluSx3_exp_ +/-) or both (AluSx3_exp_ -/-) alleles confirming successful editing of both lines.

**D. CNV** analysis of isogenic lines with QDNAseq v1.42.0. The scatter plots show the log2 copy-number variation for a 100kb binning, based on local coverage, corrected by mappability and CG content. The effective estimated CNV number, represented by the orange horizontal segments, never reaches the threshold values for CNV=3 or CNV=1 (0.58 and - 1.0, respectively, corresponding to log2(3/2) and log2(1/2)), indicating that the isogenic lines have overall a normal CNV profile.

**Supplementary Figure 5. Morphological assessment and expression analysis in iPSC-sensory neurons**

**A.** Quantification of **neurite** outgrowth as ratio of neurite area to number of nuclei in sensory neurons. Data are shown as single data points, mean values and standard error from at least two different experiments. The statistical significance has been calculated using the Brown-Forsythe and Welch ANOVA with Dunnet’s T3 multiple comparisons test. Scale bar = 50 uM

**B. Uncorrected** (AluSx3_exp_ +/+) and corrected (AluSx3_exp_ +/- and AluSx3_exp_ -/-) iPSC lines from two CANVAS patients were analysed by RNA-seq in their undifferentiated stage (day 0) or after sensory neuronal differentiation (day7). Principal Component Analysis (PCA) for all expressed genes are shown. Left-most plot shows that samples cluster primarily according to their differentiated state along the PC1 axis, and secondarily by patient lineage along PC2 axis. Middle plot shows batch effects in neuronal differentiation. Right-most plot shows sensory neurons from corrected lines situated lower on the PC2 axis than their matched uncorrected line for each lineage-batch combination. Technical replicates n=2 for undifferentiated iPSC and n=3 and n=2 for CANVAS1 and CANVAS2 day7 sensory neurons, respectively.

**C.** Heat map of RNA-seq expression of sensory neuronal marker genes in undifferentiated (iPSCs) and day 7 sensory neurons for all lines and differentiation batches.

### Supplementary Figure 6. Transcriptional changes in CANVAS iPSC sensory neurons

**Volcano** plot (**top**) and Gene Ontology (GO) analysis (**bottom**) of differentially expressed genes in CANVAS uncorrected (AluSx3_exp_ +/+) vs all corrected (AluSx3_exp_ +/- and AluSx3_exp_-/-) iPSC-derived sensory neurons. N=3 technical replicates for CANVAS1 and n=2 technical replicates for CANVAS2. The five most significant GO pathways are shown.

**B.** Ratio of Normalised read counts for all subunits of Replication Factor Complex (*RFC1-5)* in heterozygous (AluSx3_exp_ +/-) and homozygous (AluSx3_exp_ -/-) isogenic control iPSC-derived sensory neurons (day 7) compared to their unedited CANVAS (AluSx3_exp_ +/+) parent line. Technical replicates N=3 for CANVAS1 and n=2 for CANVAS2 lines, data are shown single data points, mean and standard error. The statistical significance was calculated using paired t test analysis.

**C.** DEXSeq plots of *RFC1* splicing showing similar splicing and alternative exon usage in unedited CANVAS (AluSx3_exp_ +/+), heterozygous (AluSx3_exp_ +/-) and homozygous (AluSx3_exp_ -/-) corrected isogenic control iPSC sensory neurons. Technical replicates N=3 for CANVAS1 and n=2 for CANVAS2 lines.

**Supplementary Figure 7. Morphological assessments of iPSC-motor neurons**

**A.** Quantification of neurite outgrowth as ratio of neurite area to number of nuclei in motor neurons. Data are shown as single data points, mean values and standard error from at least two different experiments. The statistical significance has been calculated using unpaired t-test.

**B.** Representative immunofluorescence of post-mitotic iPSC-induced lower motor neurons showing robust staining for lower-motor neuron marker HB9 (or MNX1) and neuronal markers Microtubule Associated Protein 2 (MAP2), and β3-Tubulin (TUBB3).

***Supplementary Figure* 8. PCR-amplicon based targeted long-read sequencing of *RFC1* in iPSC sensory and motor neurons**

**A.** Full-length transcripts identified in at least one sample. Novel transcripts are named BambuTx*N.*

**B.** Heatmap showing expression level of each transcript for all samples. Expression is measured as log-transformed Counts Per Million (number of reads normalized by sample sequencing depth).

**Supplementary Fig. 9. Knockdown of RFC1 in iPSC-derived sensory neurons**

**A.** Day 14 sensory neurons were transduced with either RFC1 or scramble shRNA lentivirus. The relative number of surviving neurons was tracked using live-cell imaging every 8 hours in an Incucyte S3 microscope. The dashed line represents the expected start of RFC1 knockdown. The experiment was carried out in triplicates.

**B.** Quantitative RT-PCR for *RFC1* expression in day 14 iPSC-derived sensory neurons treated with either RFC1 or scramble shRNA. Relative expression levels (RQ) of *RFC1* are normalized to *TBP* using the 2^–ΔΔCt method, showing a 83% reduction of *RFC1* in RFC1 shRNA treated lines compared to scramble shRNA.

**Supplementary Figure 10. CANVAS lymphoblastoid cell lines show increased sensitivity to platinum-based drugs**

**A.** Lymphoblastoid cell lines from CANVAS patients (n=3) and healthy controls (n=3) were treated for 24h and 48h with different concentrations of Oxaliplatin and cell viability was evaluated. Data are shown as single data points, mean and standard error. Unpaired t-test with Welch correction was performed to compare two groups assuming unequal variance between the two. Experiments were carried out on 5 CANVAS and 5 control LCLs and each condition was replicated at least three times.

**B.** Western blot analysis of the expression of cleaved Caspase-3 and γH2A.X factors in cells from CANVAS patients (P) and healthy controls (C) grown in the presence of 5 μM of oxaliplatin for 24 h (left) and 48 h (right). Statistical significance was calculated using Brown-Forsythe and Welch ANOVA test with Dunnett’s T3 multiple comparisons post-hoc test.

**C.** Percentages of lymphoblastoid cell lines (LCLs) from CANVAS patients and healthy individuals in different cell cycle phases (sub-G1, G0/G1, S, G2) exposed to 30 μM cisplatin for 1h and then recovered in complete medium for 0 h (left), 24 h (middle) and 48 h (right). Mean values ± standard deviation of measurements from two different experiments (n=2) are shown.

**D.** Percentage of sub-G1 fraction of cells from healthy controls (black) and CANVAS patients (violet). Mean values ± standard deviation of measurements from two different experiments (n=2) are shown. Over the bars is reported the fold increase in sub-G1 fraction of 30 μM cisplatin treated cells compared to untreated ones.

**E.** Bar graph representing the proportion of alive, early apoptotic, late apoptotic/necrotic cells after 24 h recovery in the presence of 5 μM or 10 μM cisplatin. Mean values ± standard deviation of measurements from two different experiments (n=2) are shown. Over the bars is reported the fold increase in apoptotic or late apoptotic/necrotic treated cells compared to untreated ones.

**F.** Western blot analysis of cleaved Caspase-3 in CANVAS patients (P) and healthy controls (C) after cisplatin pulse (1h) and recovery (24h). Statistical significance was calculated using Brown-Forsythe and Welch ANOVA test with Dunnett’s T3 multiple comparisons post-hoc test.

**Supplementary Figure 11. Generation and phenotyping of *Gnf1* (RFC1 orthologue) Drosophila melanogaster model.**

**A.** Schematic of the *Gnf1* locus. Exons – white blocks; untranslated regions (UTRs) – grey blocks; introns – line. Arrow shows exonic region targeted by *Gnf1* shRNA.

**B.** Quantitative PCR validation of *Gnf1* knockdown in dissected adult male fly brains. n=3 independent replicates per genotype P value indicated as * (p<0.05) acquired by unpaired t test with Welch’s correction.

**C.** Visualisation of single-cell RNA-seq data from the adult *Drosophila* central nervous system using SCOPE (Davie et al. 2018). *Gnf1* expression (red) is compared to the post-mitotic neuronal marker *nsyb* (blue), illustrating widespread expression of Gnf1 in neurons.

**E.** Schematic of the *Drosophila* Activity Monitor (DAM) system. Flies housed individually in glass tubes are shown crossing a central infrared beam, allowing the number of beams breaks over time to be used as a readout of changes in locomotor activity.

**E-G**. Patterns of locomotor activity in adult male flies of differing ages, expressing Gnf1 shRNA in post-mitotic neurons (*nsyb > gnf1* shRNA) and driver/shRNA alone controls. White bar: lights on; black bar: lights off. **E**: 3-7 days old flies (n=9-18); **F**: 21-23 days old flies (n=19-60); and **G**: 40-42 days old flies (n=15-23).

**Supplementary Figure 12. Effects of cisplatin treatment on flies expressing Gnf1 shRNA in post-mitotic neurons (*nsyb > gnf1* shRNA) and driver/shRNA alone controls.**

**A.** Survival curve showing the percentage survival of *nsyb > gnf1* shRNA flies compared to controls in response to cisplatin treatment. n=90-100 per genotype.

**B.** Patterns of locomotor activity over a 12 h light: 12 h dark period in control and neuronal Gnf1 knockdown 12 day old adult male flies on cisplatin treatment. Note the prominent advance of anticipatory increases in movement prior to lights-off in the Gnf1 knockdown background compared to controls. White bar: lights on; black bar: lights off.

**C-D**. Locomotor activity in control and neuronal Gnf1 knockdown 12 day old adult male flies on cisplatin treatment (quantified as beam breaks in the *Drosophila* Activity Monitor) during two 2 h period: ZT9-11 (prior to lights-off) (**C**) or ZT12-14 (after lights-off) (**D**). Error bars in **C-D**: standard error of the mean (SEM). Central line in dot plots: mean. P values are indicated, acquired via Mantel-Cox Log rank test (a) and Kruskal-Wallis test with Dunn’s T3 multiple comparisons post-hoc test (c, d)

**Supplementary Figure 13. Recursive Directional ligation**

A range of (CCCTT)_n_ and (CTTTT)_n_ containing plasmids were generated through recursive directional ligation (McDaniel 2010). Oligonucleotides were appropriately designed to cointain (CCCTT)_12_- or (CTTTT)_12_ repeated sequence flanked by Esp3I and PaqCI recognition sites. Additionally, they contained a 5’ NotI recognition site and a 3’ BamHI recognition sites, also present in pBluescript SK+ plasmid. After pBluescript SK+ digestion with NotI and BamHI restriction enzymes (A), repeat containing oligonucleotides – in red –were annealed and inserted into the plasmid (B). Two separate double digestion reactions with PaqCI and BsaHI (C) and Esp3I and BsaHI (D) restriction enzymes released then two fragments containing the repeated sequence, with complementary overhangs. After separation through gel electrophoresis (not shown), ligation of the two fragments yielded a plasmid with seamlessly joined repeat units into a sequence double in size (E). Repetition of the same process yielded a controlled expansion of the repeated sequence within the plasmid.

Measurement of exact repeat size of different vectors generated is summarized in

**Supplementary Table 4**.

## SUPPLEMENTARY TABLES

**Supplementary Table 1**. Demographic and clinical data of brain donors and patient-derived cell lines

**Supplementary Table 2.** Differently expressed genes from RNA-seq from Cerebellar Hemisphere and Frontal Cortex

**Supplementary Table 3**. Variation of *RFC1* expression across tissues between AAGGG carriers and non-expanded individuals as part of GTEx project.

**Supplementary Table 4.** Differently expressed genes from RNA-seq from iPSC sensory neurons

**Supplementary Table 5.** Repeat insert size in pBluescript, luciferase and pGINT vectors.

|  | Measured repeat number (LRS) |  |  |  |  |  |
| --- | --- | --- | --- | --- | --- | --- |
|  | Firefly luciferase vector |  | pGINT EGFP splicing vector |  | pBluescript vector |  |
| Expected repeat number | CCCTT | CTTTT | CCCTT | CTTTT | CCCTT | CTTTT |
| 12x | 13 | 12 | 13 | 12 | 12 | 12 |
| 24x | 24 | 23 | 22 | 23 | 22 | 23 |
| 48x | 35 | 47 | 36 | 45 | 36 | 45 |
| 60x | 68 | 60 | 68 | 56 | 68 | 56 |

**Supplementary Table 6:**
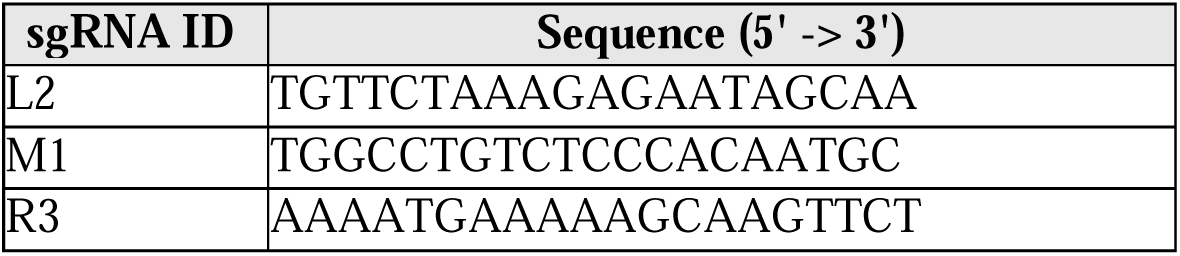
Guide RNA Sequences for iPSC editing.

**Supplementary Table 7:**
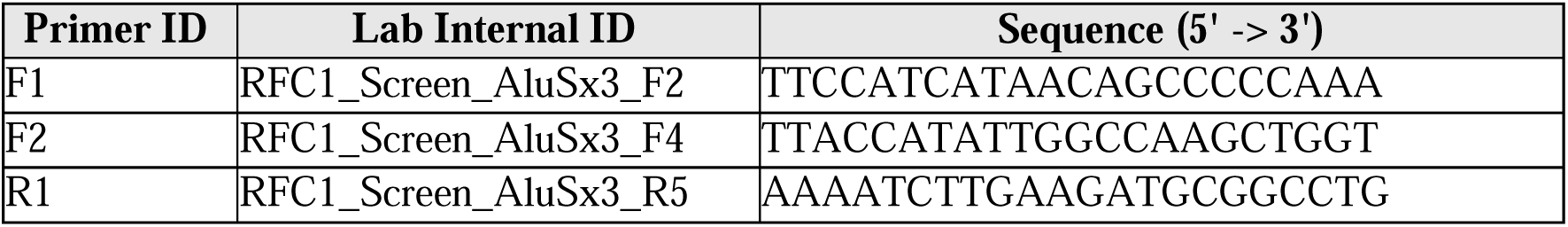
Primer sequences used to screen iPSC clones after editing.

## Notes

### Competing Interest Statement

The authors have declared no competing interest.

### Funding

- European Research Council: 101165557
- National Ataxia Foundation
- Medical Research Council: MR/T001712/1
- Fondazione Cariplo: 2019-1836
- Charcot-Marie-Tooth Association: SR-202504
- AFM-Telethon: 28813
- Fondazione Regionale per la Ricerca Biomedica: 1751723
- Muscular Dystrophy UK: 24GRO-PG24-0719-1
- Guarantors of Brain

