## Supplementary figures and images for "CANVAS causing AAGGG repeat expansions cause tissue-specific reduction in RFC1 expression and increase sensitivity to DNA damage"

### Suppl Fig 1.tif

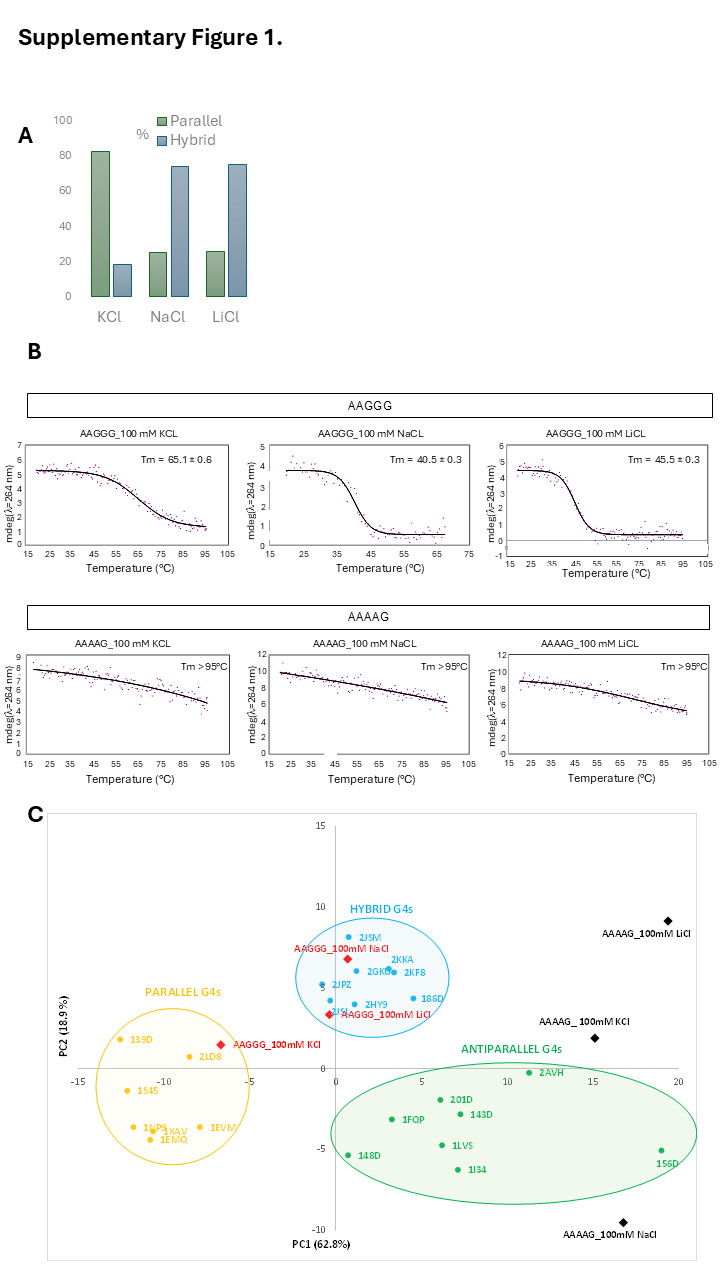

### Suppl Fig 2.tif

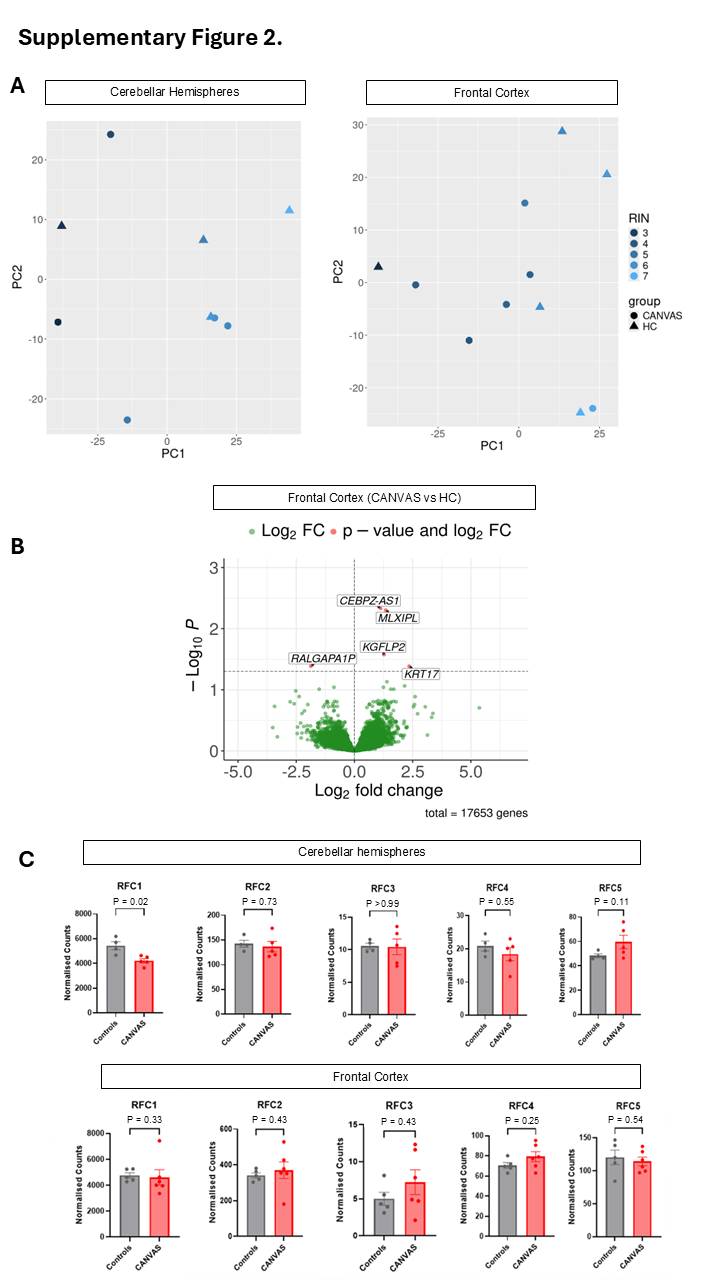

### Suppl Fig 2D.tif

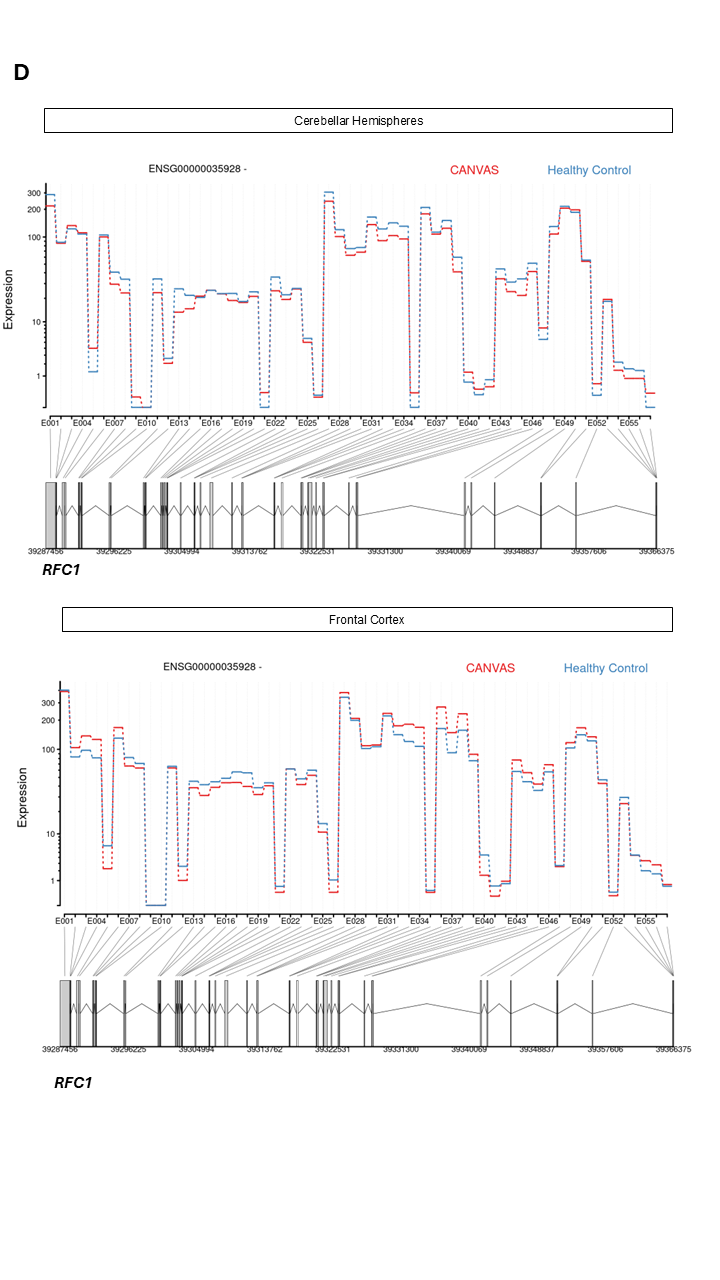

### Suppl Fig 3.tif

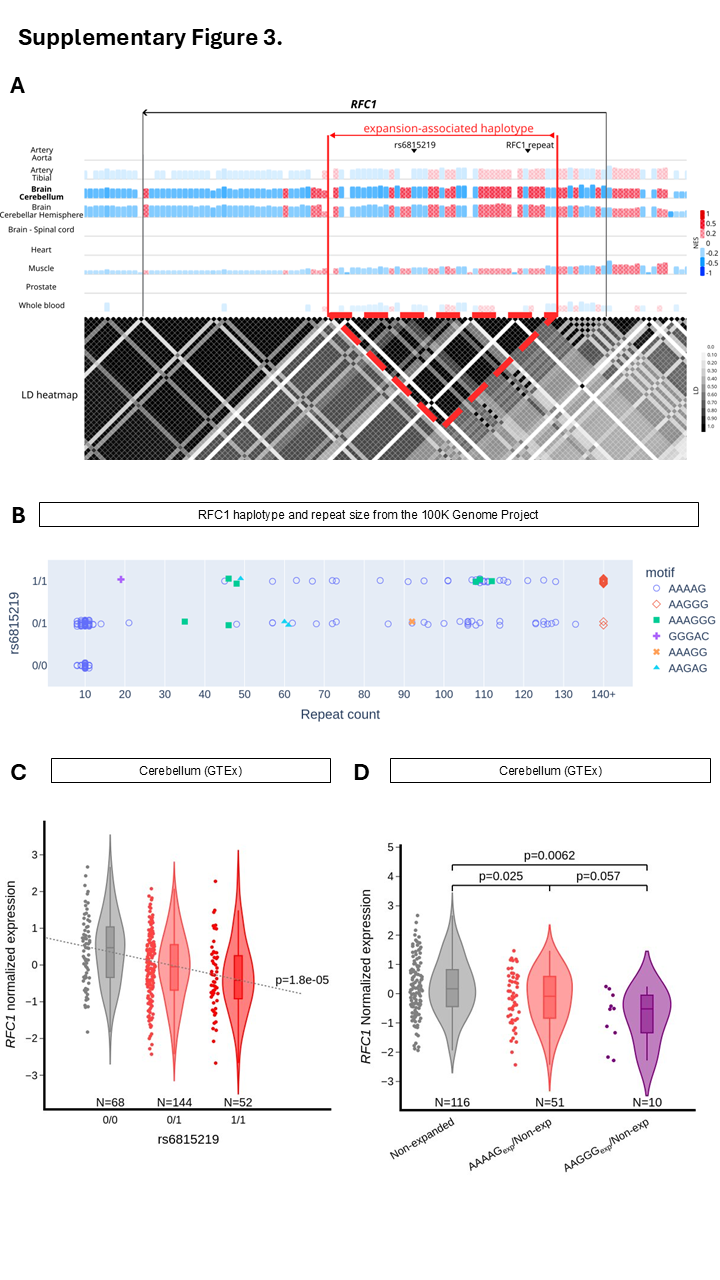

### Suppl Fig 4A.tif

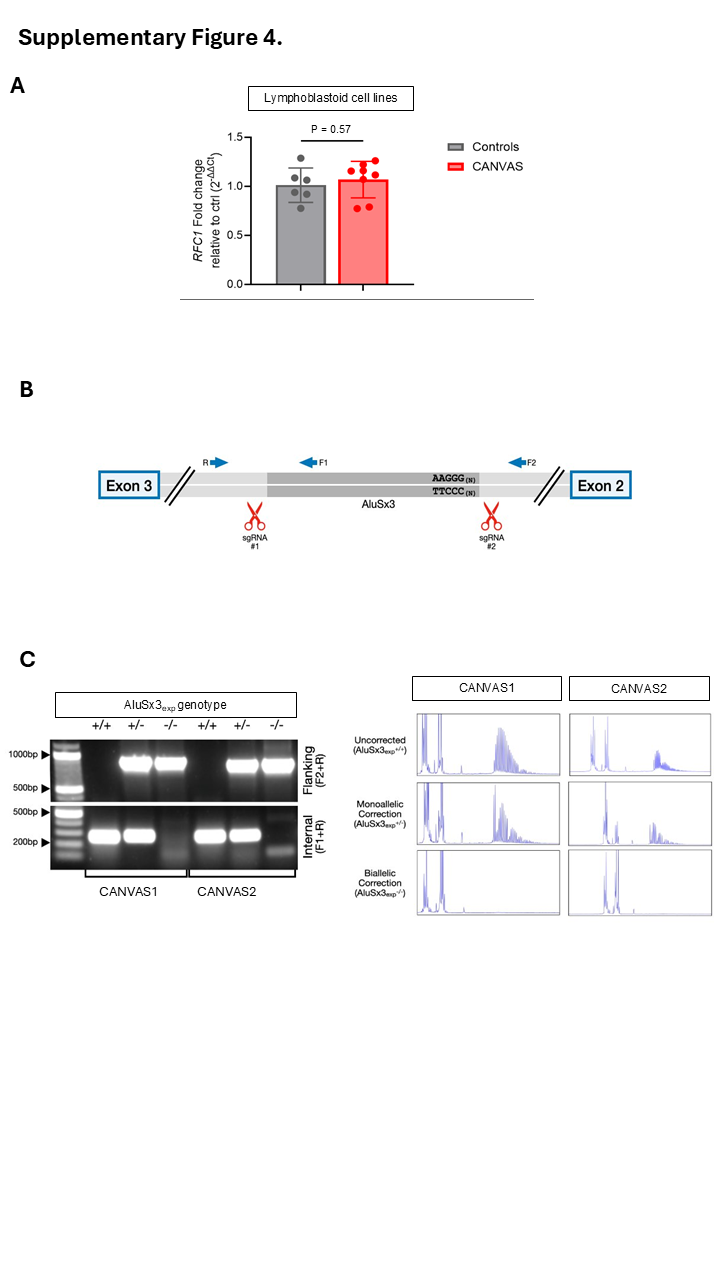

### Suppl Fig 4D.tif

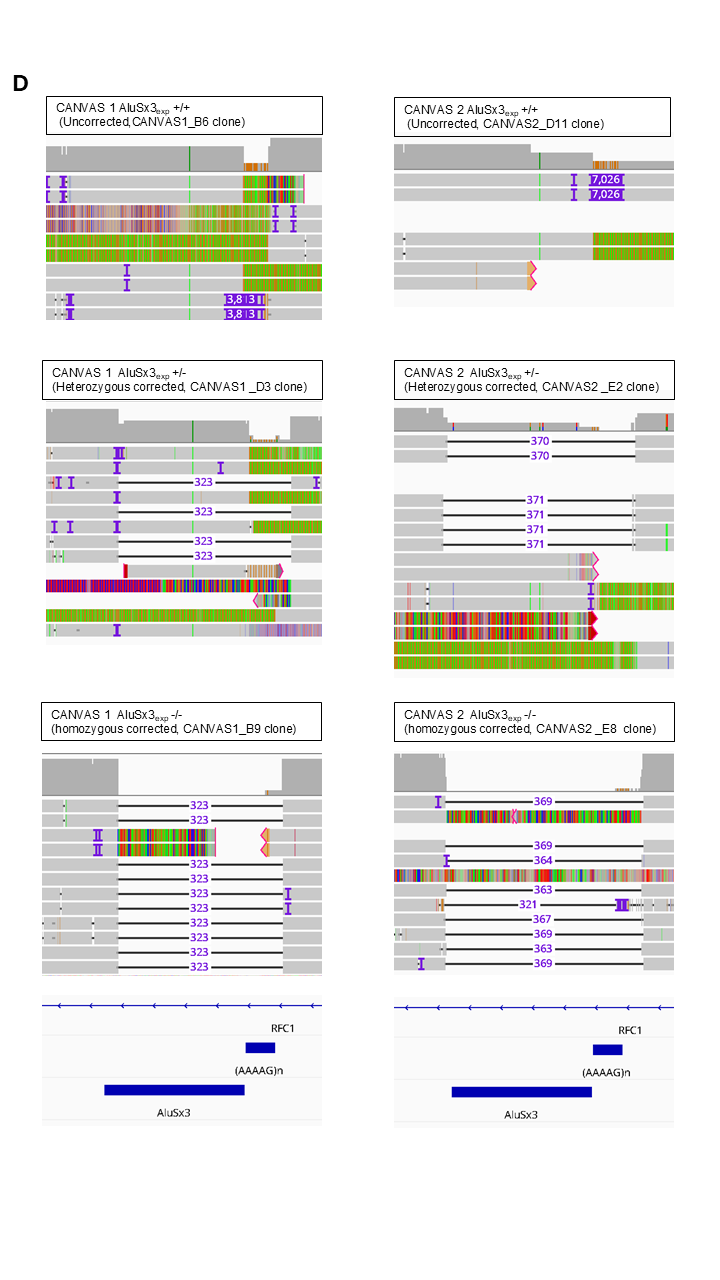

### Suppl Fig 4E.tif

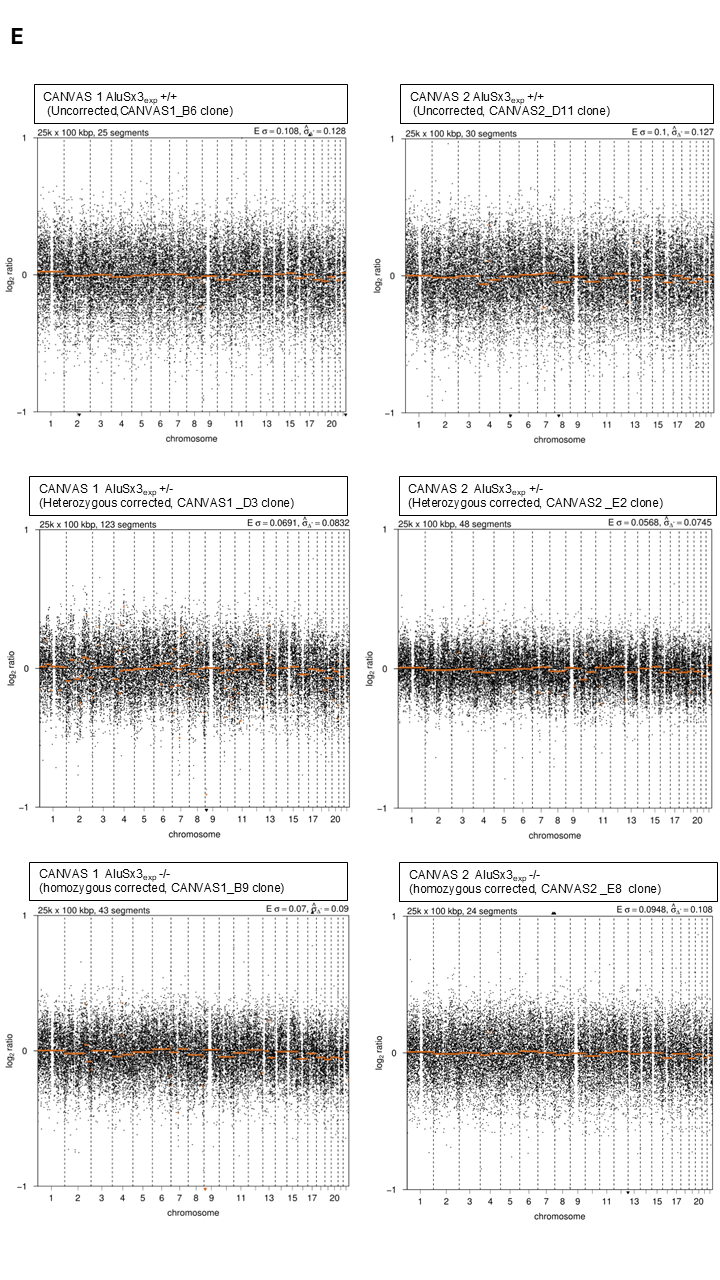

### Suppl Fig 5.tif

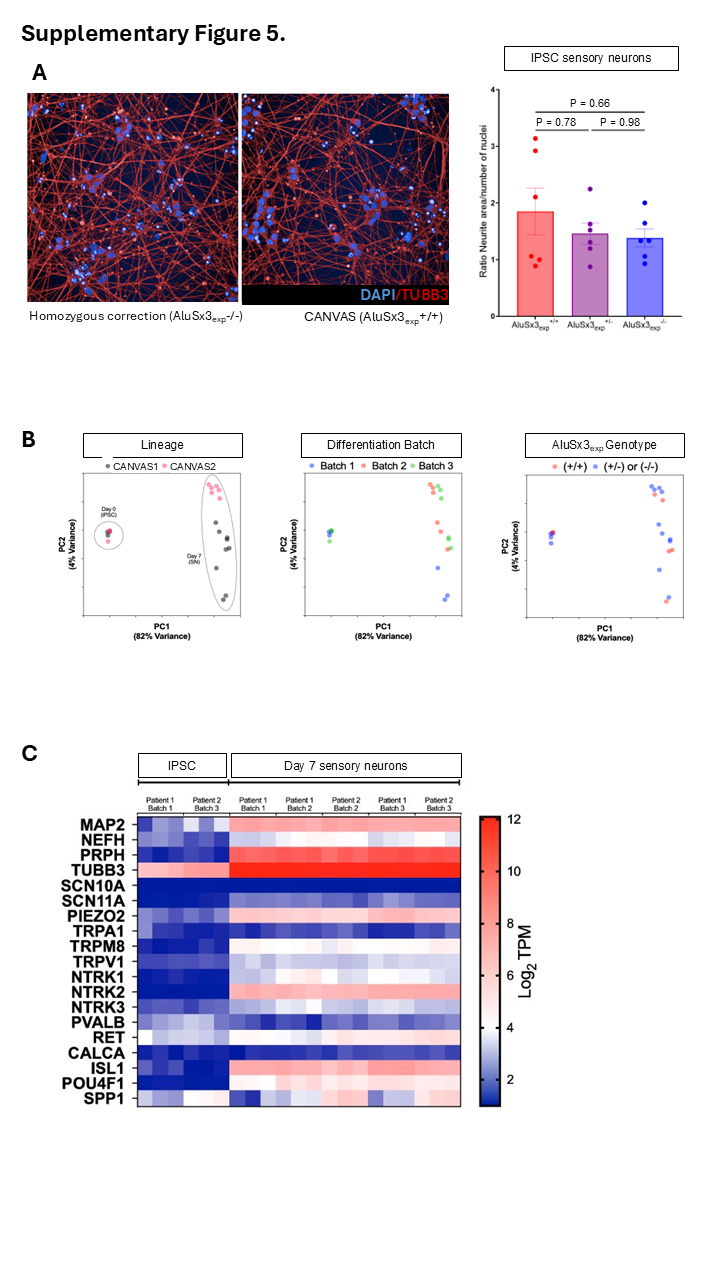

### Suppl Fig 6.tif

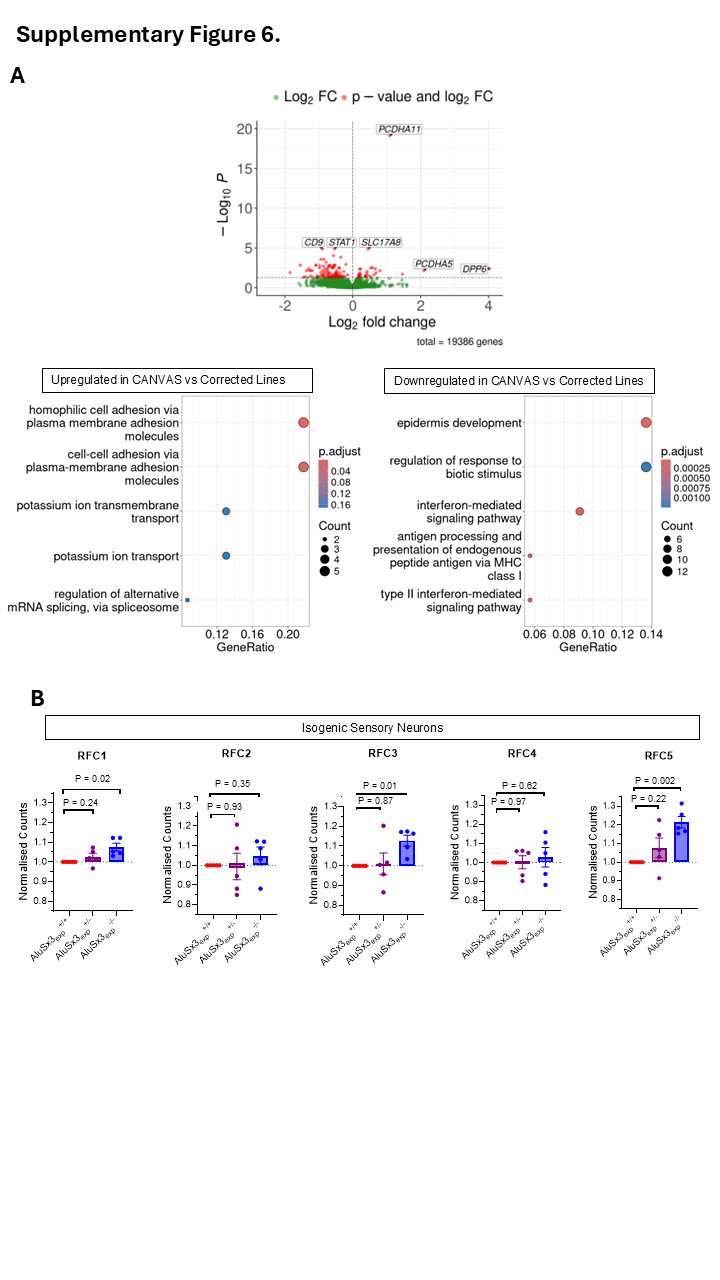

### Suppl Fig 7.tif

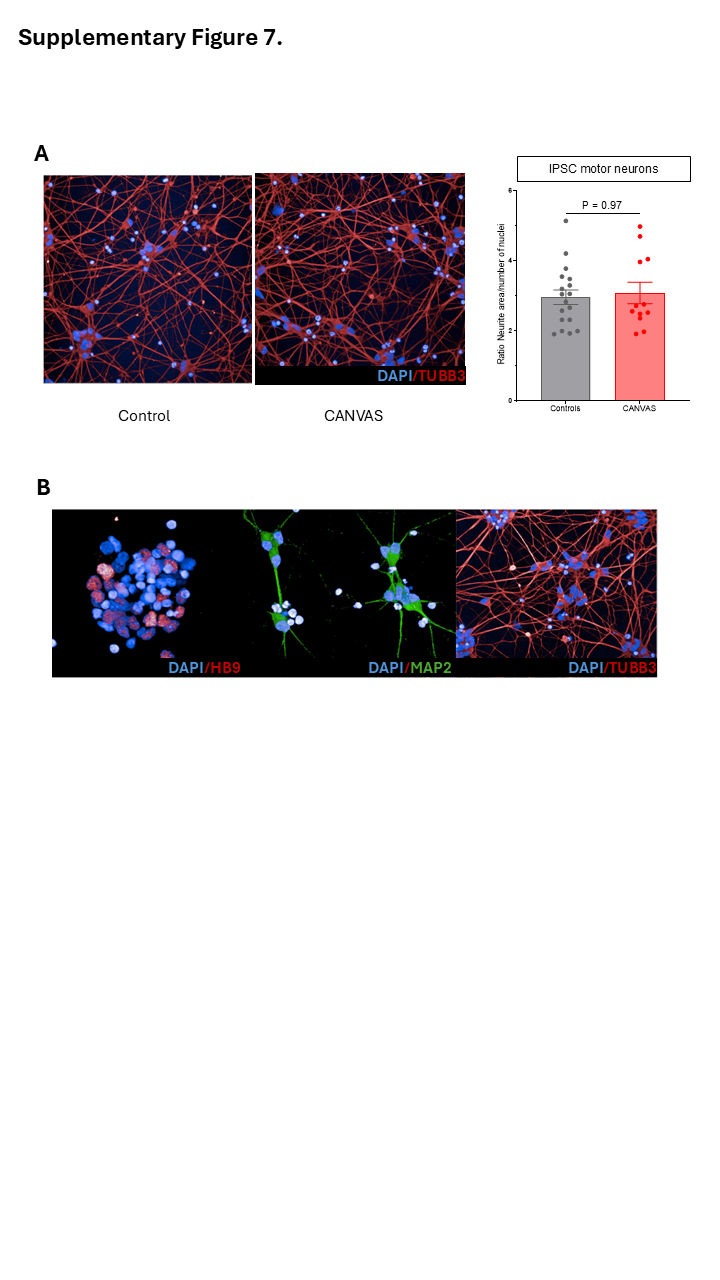

### Suppl Fig 8.tif

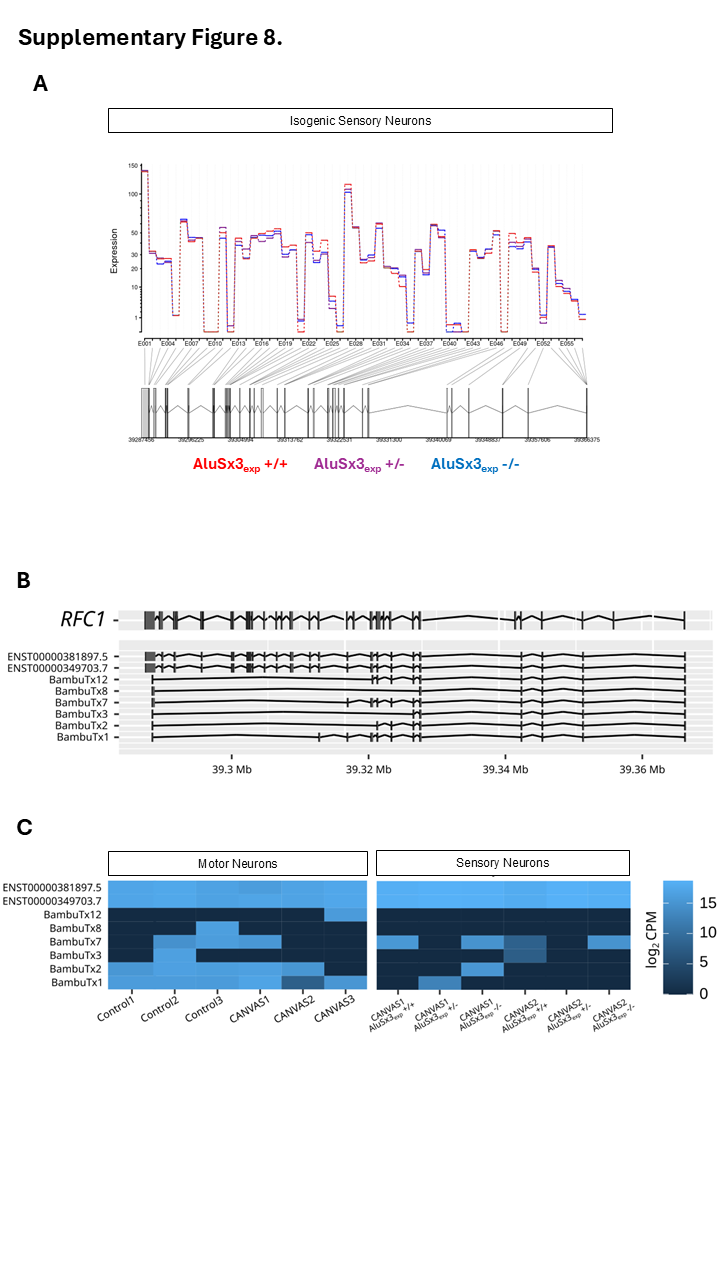

### Suppl Fig 9.TIF

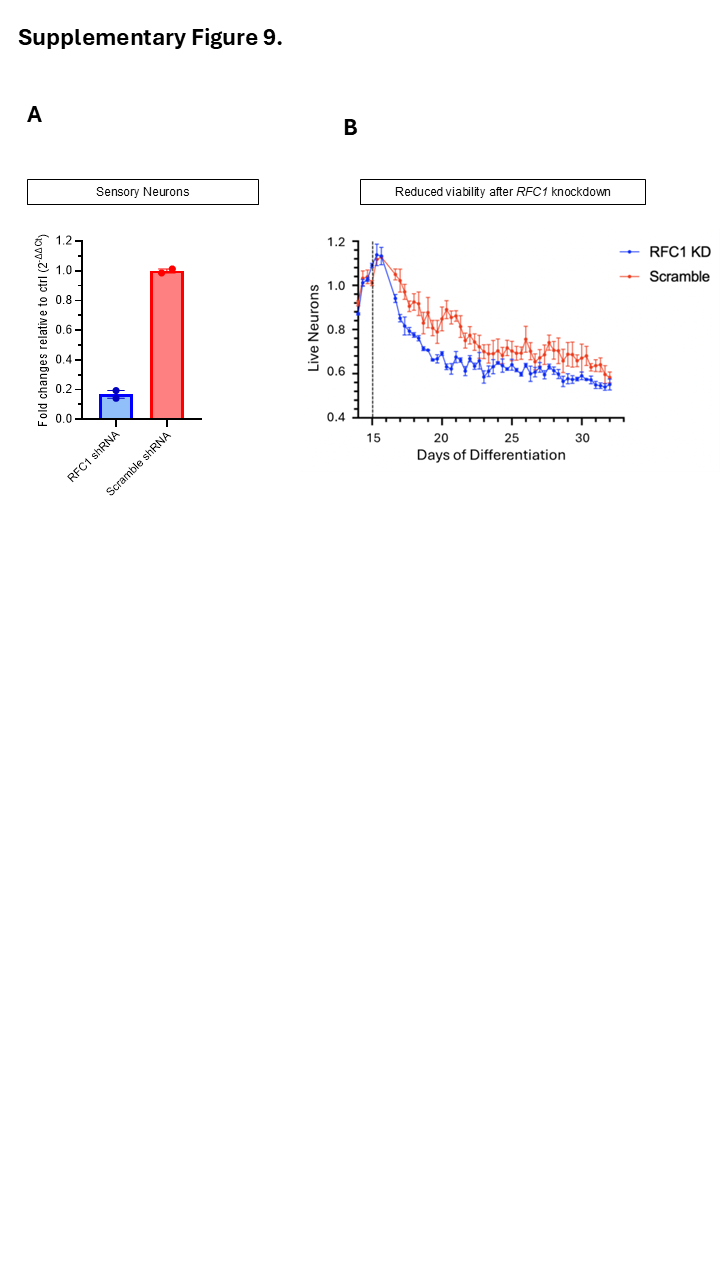

### Suppl Fig 10.TIF

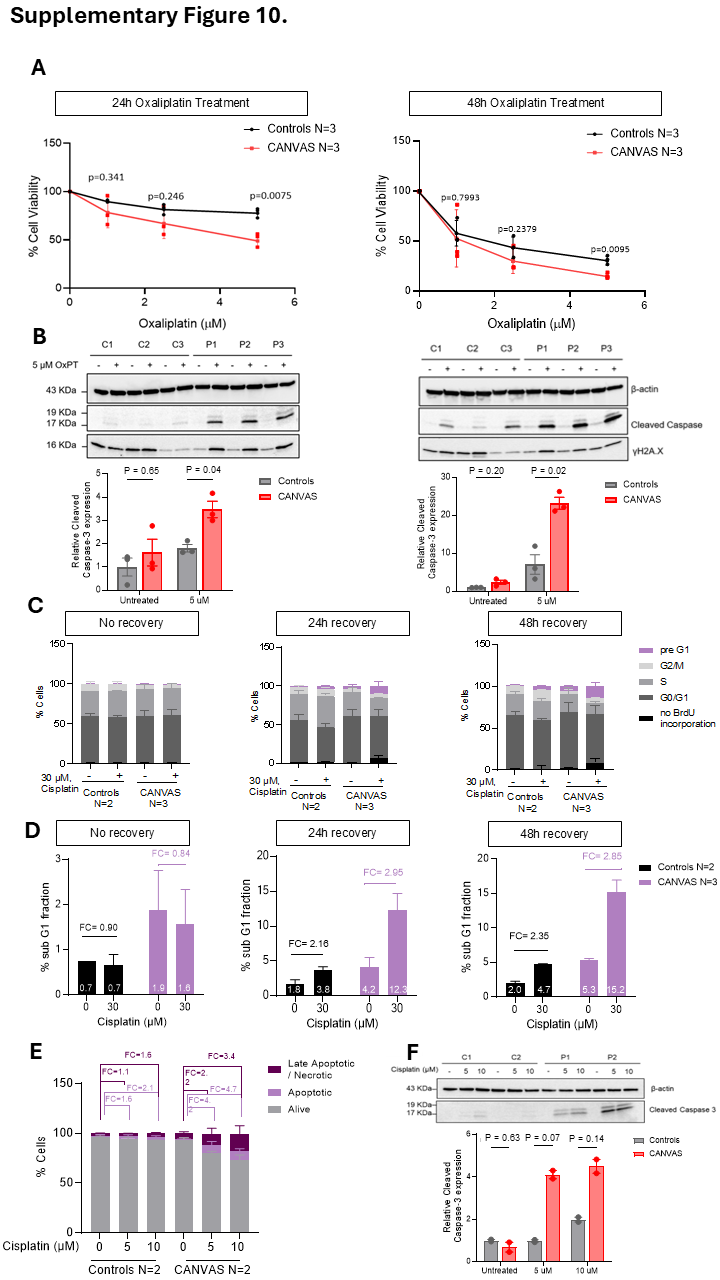

### Suppl Fig 11.TIF

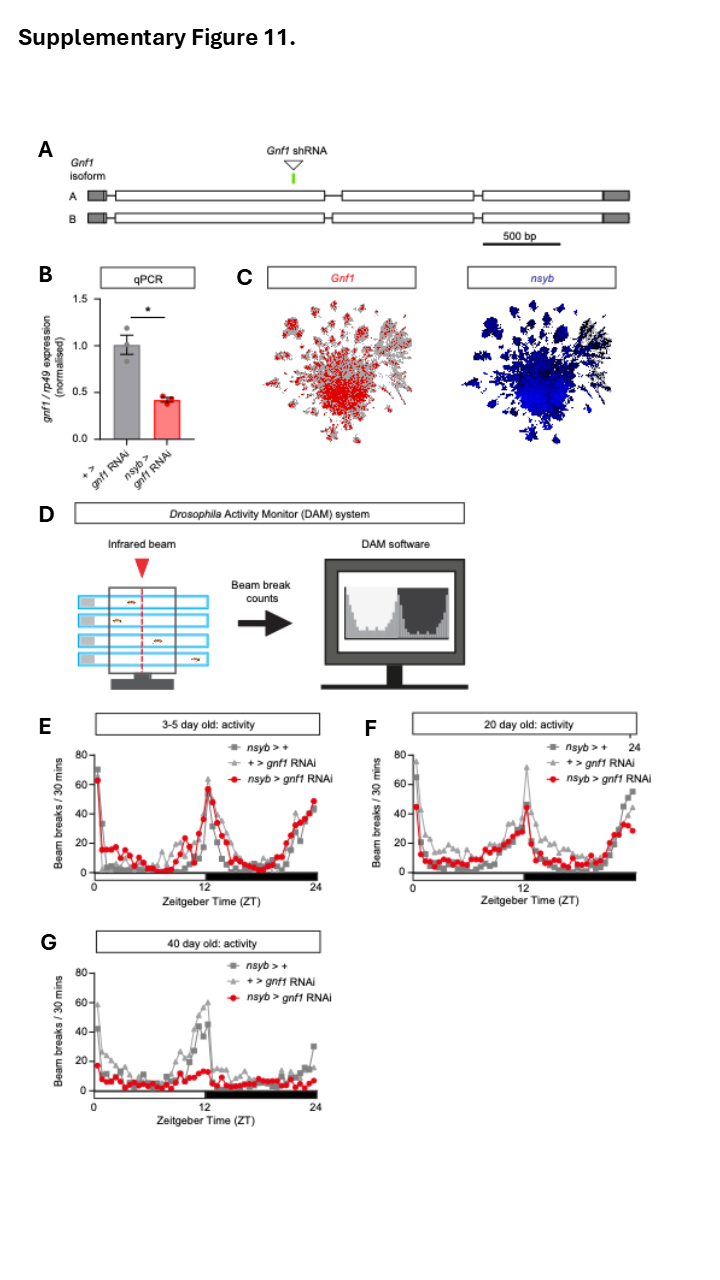

### Suppl Fig 12.TIF

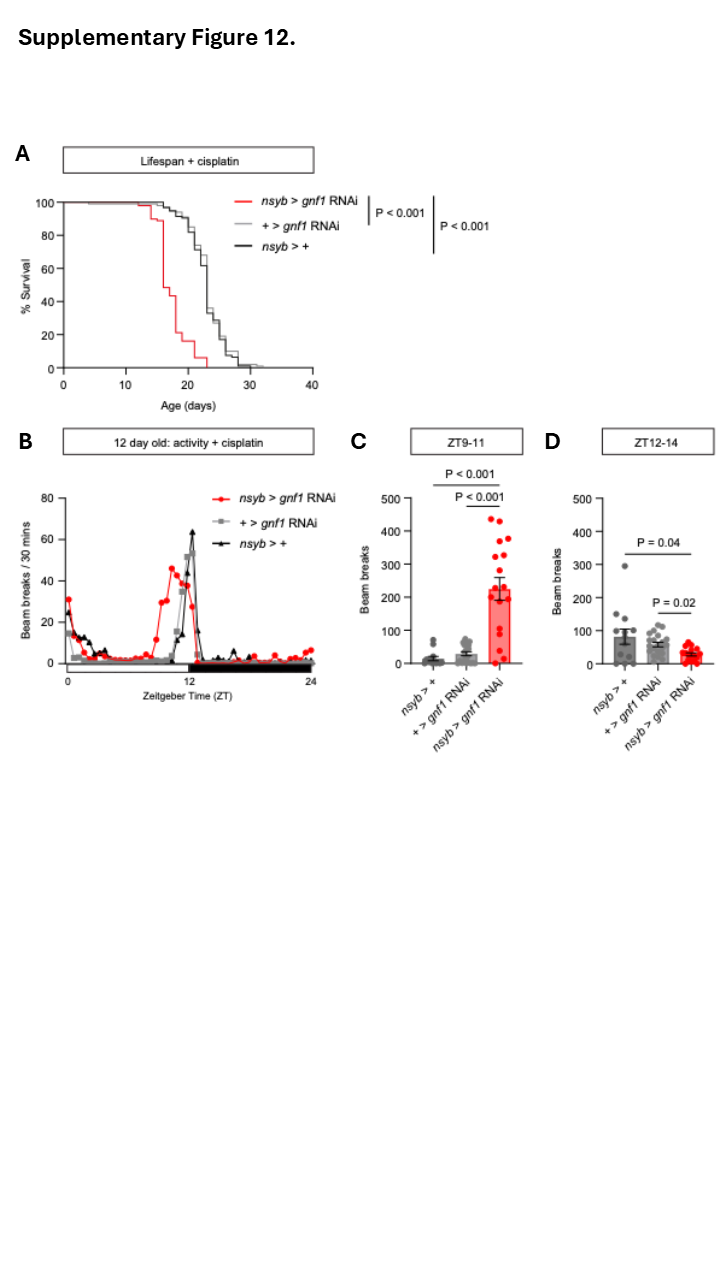

### Suppl Fig 13.TIF

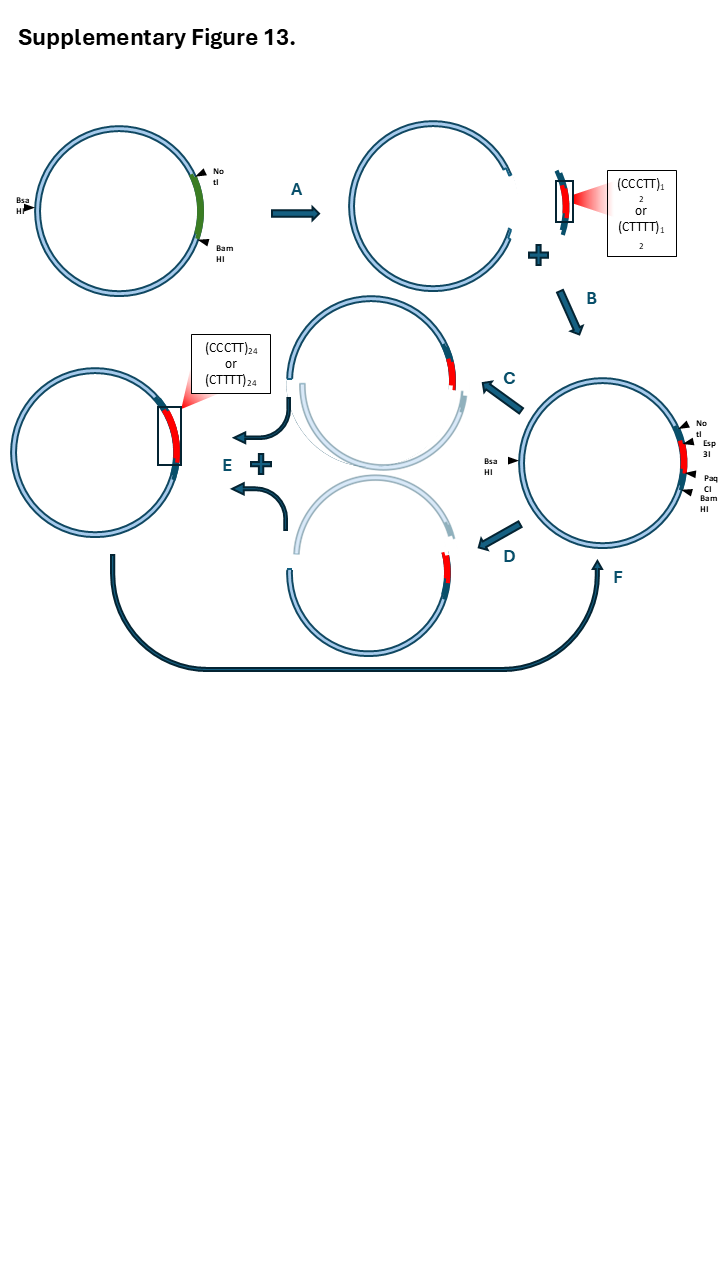
